# BAP1 activity regulates PcG occupancy and global chromatin condensation counteracting diffuse PCGF3/5-dependent H2AK119ub1 deposition

**DOI:** 10.1101/2020.12.10.419309

**Authors:** Eric Conway, Federico Rossi, Simone Tamburri, Eleonora Ponzo, Karin Johanna Ferrari, Marika Zanotti, Daniel Fernandez-Perez, Daria Manganaro, Simona Rodighiero, Diego Pasini

**Author notes:** Correspondence (D.P.). **Contact information of the corresponding author/lead contact** Diego Pasini, Ph.D., Department of Experimental Oncology, European Institute of Oncology, Via Adamello, 16. 20139, Milano, Italy.

## Abstract

BAP1 is recurrently mutated or deleted in a large number of diverse cancer types, including mesothelioma, uveal melanoma and hepatocellular cholangiocarcinoma. BAP1 is the catalytic subunit of the Polycomb Repressive De-Ubiquitination complex (PR-DUB) which removes PRC1 mediated H2AK119ub1. We and others have shown that H2AK119ub1 is essential for maintaining transcriptional repression and contributes to PRC2 chromatin recruitment. However, the precise relationship between BAP1 and PRC1 remains mechanistically elusive. Using embryonic stem cells, we show that a major function of BAP1 is to restrict H2AK119ub1 deposition to target sites. This increases the stability of PcG complexes with their targets and prevents diffuse accumulation of H2AK119ub1 and H3K27me3 modifications. Loss of BAP1 results in a broad increase in H2AK119ub1 levels that are primarily dependent on PCGF3/5-PRC1 complexes with a mechanism that is reminiscent of X-chromosome inactivation. Increased genome-wide H2AK119ub1 levels titrates away PRC2 from its targets and stimulates diffuse H3K27me3 accumulation across the genome. This decreases the activity of PcG repressive machineries at physiological targets and induces a general compaction of the entire chromatin. Our findings provide evidences for a unifying model that resolves the apparent contradiction between BAP1 catalytic activity and its role *in vivo*, uncovering molecular vulnerabilities that could be useful for BAP1-related pathologies.

## Introduction

BAP1 is a ubiquitin C-terminal hydroxylase (UCH) enzyme with an affinity for Histone H2A lysine 119 mono-ubiquitination (H2AK119ub1) as a substrate (Sahtoe et al., 2016; Scheuermann et al., 2010). It’s recurrently mutated or deleted with particularly high frequency in mesothelioma (~36%), uveal melanoma (~40%), hepatocellular cholangiocarcinoma (~26%) and renal clear cell carcinoma (~24%) among others, delineating tumor suppressive function of BAP1 in these cancers (Carbone et al., 2013; Cerami et al., 2012; Gao et al., 2013). While these mutations are primarily somatic, familial mutations in BAP1 have been reported to predispose patients to mesothelioma and uveal melanoma (Testa et al., 2011). BAP1 forms a multi-subunit chromatin modifying complex, called Polycomb Repressive De-ubiquitinase complex (PR-DUB), which contains the ASXL1-3 proteins along with other subunits FOXK1/2, HCFC1, OGT, MBD5/6 and KDM1B (Hauri et al., 2016; Kloet et al., 2016). ASXL1-3 are essential for DUB catalytic activity through formation of a dimer with BAP1 (Sahtoe et al., 2016; Scheuermann et al., 2010). The ASXL subunits are also mutated in myeloid disorders and neurodevelopmental disorders (Bohring-Opitz and Bainbridge-Ropers syndromes) tightly linking the PR-DUB complex and H2AK119ub1 activity to disease pathogenesis (Bainbridge et al., 2013; Bejar et al., 2011; Gelsi-Boyer et al., 2009; Hoischen et al., 2011).

Although the clinical relevance of BAP1 to these cancers has been well established, the precise mechanism of BAP1 in tumor suppression remains unclear. The DUB activity of BAP1 for H2AK119ub1 suggests an antagonistic relationship between BAP1 and the PRC1 complex, which contains the RING1A and RING1B E3 ligases that catalyse H2AK119ub1 (Blackledge et al., 2015). PRC1 has vital roles in transcriptional repression, particularly during development and cell fate decisions (Deevy and Bracken, 2019). It is comprised of six major complex subtypes, each defined by the PCGF1-6 paralogs that forms the core catalytic heterodimer with RING1A/B (Gao et al., 2012). These sub-complexes can be broadly divided into canonical (PCGF2/4-containing) or non-canonical (PCGF1/3/5/6-containing) PRC1 with specific genomic localisations and catalytic activities (Fursova et al., 2019; Scelfo et al., 2019). Recently, we and others have shown that PRC1 mediated H2AK119ub1 is required for the repression of PRC1 target genes in embryonic stem cells (ESCs) and proper recruitment of PRC2 complexes to these sites (Blackledge et al., 2020; Tamburri et al., 2020).

The PRC2 complex is responsible for the generation of the repressive H3K27me3 modification through the catalytic EZH1/2 subunits of the complex (Ferrari et al., 2014; Lavarone et al., 2019). Like PRC1, there are a number of complex subtypes that are broadly divided into PRC2.1 (containing PCL1-3 with either EPOP or PALI1) or PRC2.2 (containing JARID2 and AEBP2) (Beringer et al., 2016; Conway et al., 2018; Grijzenhout et al., 2016; Hauri et al., 2016). Recently, the mechanisms of PRC2 complex recruitment have been explored highlighting divergent recruitment models for the sub-complexes, with each auxiliary subunit having specific genomic targeting properties for the catalytic PRC2 core formed by EZH1/2, SUZ12 and EED (Glancy et al., 2021; Healy et al., 2019; Højfeldt et al., 2019). While PRC2.1 is primarily recruited at target sites via the CpG island recognition capacity of PCL1-3 (Choi et al., 2017; Healy et al., 2019; Li et al., 2017), PRC2.2 largely depends on PRC1 activity for chromatin occupancy (Blackledge et al., 2020; Healy et al., 2019; Tamburri et al., 2020) by affinity of its specific subunits, AEBP2 and JARID2, for H2AK119ub1 (Cooper et al., 2016; Kasinath et al., 2020). Moreover, H2AK119ub1 has been shown to be required for chromosome wide condensation during X-chromosome inactivation (Almeida et al., 2017), with a process that depends on the affinity of JARID2 for H2AK119ub1 (Cooper et al., 2016). Although the requirement for H2AK119ub1 in the regulation of somatic chromosome compaction and Polycomb body formation is less clear, (Boyle et al., 2020; Kundu et al., 2017) topological studies with complete loss of H2AK119ub1 remain to be conducted.

PR-DUB was originally discovered in Drosophila as mutation of the BAP1 homolog *Calypso* causes a classical Polycomb phenotype of anteriorisation of *Hox* gene expression, due to a loss of repression (Scheuermann et al., 2010). This gave rise to the name Polycomb Repressive De-ubiquitinase complex. However, due to the antagonism between PRC1 and PR-DUB such phenotype remains counterintuitive, with an expected Trithorax-like phenotype characterized by posteriorisation of homeotic genes expression. The confounding Drosophila phenotype is supported by *in vivo* studies in mouse where, although BAP1 knockout is lethal around gastrulation (Dey et al., 2012), mutation of either ASXL1 or ASXL2 display both Trithorax and Polycomb phenotypes in terms of homeotic transformations (Baskind et al., 2009; Fisher et al., 2010). This suggests a dual role for PR-DUB in both promoting and suppressing Polycomb activities.

While complementary knockout studies have shown that transcriptional defects in BAP1 knockout cells can be rescued by PRC1 deletion, the mechanism behind this is still unclear (Campagne et al., 2019). Curiously, some reports suggested that PR-DUB actually functions in PRC2 recruitment and that maintenance of PRC2-mediated H3K27me3 repressive modification at target genes is dependent on ASXL proteins (Abdel-Wahab et al., 2012). As a result of the mechanistic ambiguity of BAP1 function, there has been difficulty in fully characterising molecular sensitivities or synthetic lethalities for BAP1 mutated cancers (Lafave et al., 2015; Schoumacher et al., 2016).

Here we resolve these ambiguities, providing a unifying model for the role of BAP1 in both promoting and limiting PRC1 and PRC2 complex activities. We show that, while PR-DUB and PRC1/2 share very few target genes, BAP1 activity is indeed required for PRC1 and PRC2 occupancy at their target sites. Loss of BAP1 causes an indiscriminate spreading of H2AK119ub1 intergenically, which consequently titrates away PRC2 and PRC1 complexes from their target sites boosting intergenic H3K27me3 and depleting it at promoters. We show that this activity titration is dependent on the ubiquitin reading PRC2.2 subunits, AEBP2 and JARID2. In a mechanism reminiscent of X-chromosome inactivation (XCI), the intergenic spreading of Polycomb modifications causes global chromatin compaction. Like XCI, this activity is dependent primarily on the PCGF3 and PCGF5 sub-complexes of PRC1 (Almeida et al., 2017; Fursova et al., 2019). Taken together, we provide a model where BAP1 is required to maintain local concentrations of PRC2 and H3K27me3 sufficient for transcriptional regulation while exposing potential therapeutic sensitivities for BAP1 mutant cancers, such as PCGF3/5, JARID2 and AEBP2, through genetic interrogation.

## Results

### BAP1 binds active gene promoters and is excluded from PcG repressive domains

In order to investigate the relationship between PR-DUB, PRC1 and PRC2, we explored this relationship in mouse embryonic stem cells (ESCs) where the activity of Polycomb complexes have been extensively characterized (Højfeldt et al., 2019; Scelfo et al., 2019; Tamburri et al., 2020). Since BAP1 genomic occupancy has been poorly characterised until recently (Kolovos et al., 2020; Wang et al., 2018) with a dearth of commercially available ChIP grade antibodies, we generated ESC that stably expressed Flag/HA tagged BAP1 under the control of a CAG promoter (pCAG-FLAG/HA; Figure S1A). We performed ChIP-seq analyses with an HA specific antibody (Figure S1B) and identified 2291 BAP1 target genes in ESCs (Figure 1A and 1B). In addition, we also performed ChIP-seq analyses for a number of other PR-DUB complex members (ASXL1, HCFC1 and FOXK2), histone modifications (H3K27ac, H2AK119ub1 and H3K27me3) and for the PRC1 complex subunit RING1B (Figure 1A). The comparison of these ChIP-seq analyses revealed that BAP1 poorly overlapped with both PRC1 and PRC2 activities (RING1B and SUZ12). Thus, we classified target genes into two distinct groups: BAP1-only targets and RING1B-only targets (Figure 1A and 1B). Importantly, all other PR-DUB subunits accumulated specifically with BAP1-only target genes with no sign of enrichment at RING1B bound loci. Overall, these results underscore the validity of our BAP1 ChIP-seq analysis and demonstrate that BAP1 is excluded, even under conditions of ectopic expression, from PcG target loci.

**Figure 1.**
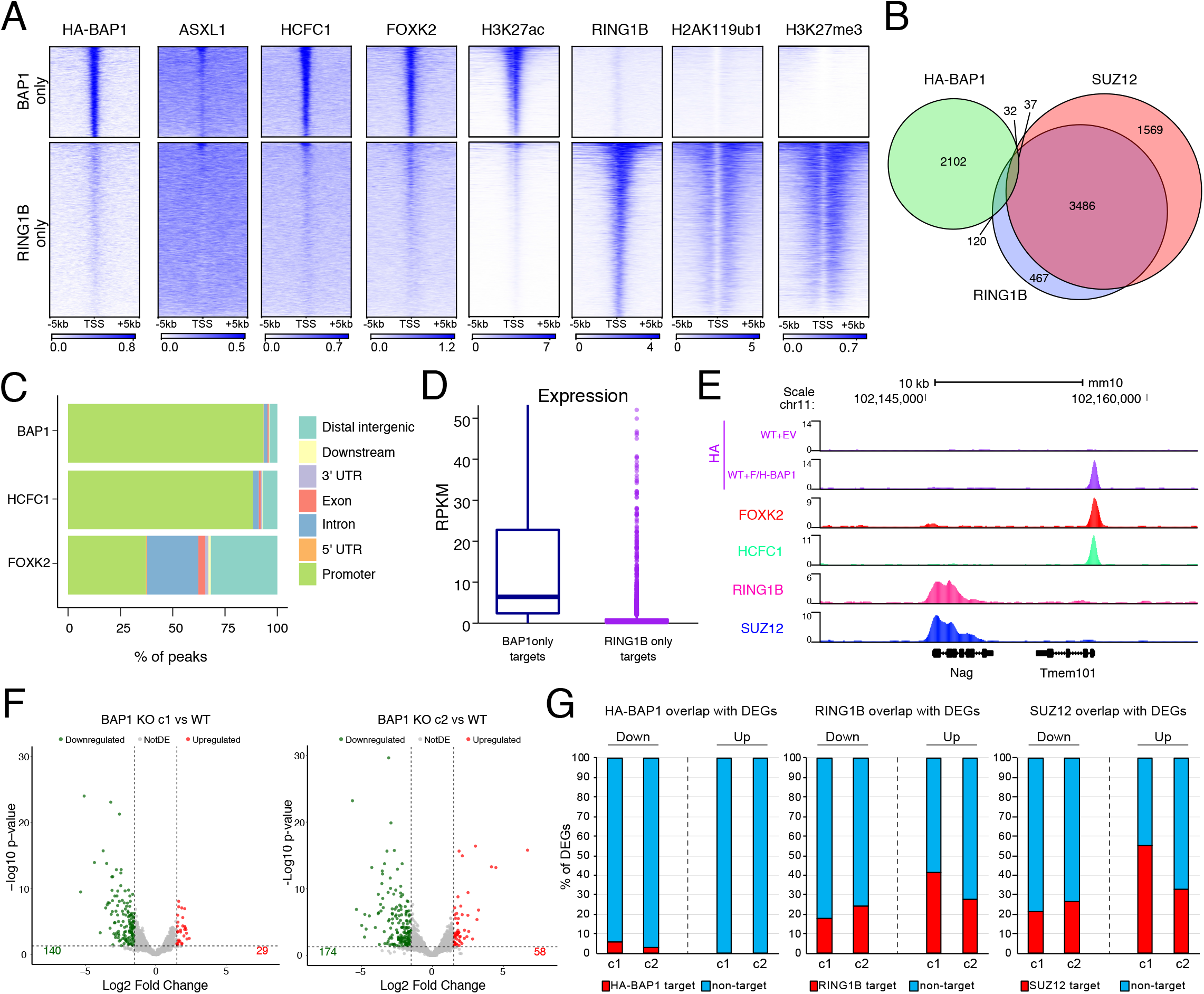
BAP1 binds active gene promoters and is excluded from PcG repressive domains. A) Heatmaps representing ChIP-seq intensity of the indicated proteins in wild-type ESC +/− 5kb of TSS. Clusters are divided into BAP1 target genes only and RING1B only targets. B) Venn diagram of HA-BAP1, SUZ12 and RING1B target genes in ESC. C) Genome-wide functional annotation of peaks generated from the indicated ChIP-seq analyses. D) Boxplots showing the expression levels obtained from RNA-seq analyses in WT mESCs for the clusters of target genes generated in A. E) UCSC genome browser snapshot of indicated ChIP-seq tracks showing an example of mutual exclusivity of PRC1/2 and PR-DUB target genes. F) Volcano plots of −log10 (p-value) against log2 fold change representing the differences in gene expression in two independent BAP1 KO ESC clones compared to WT. G) Percentage overlap of differentially expressed genes (DEG) from F with either HA-BAP1, RING1B or SUZ12 ChIP-seq targets.

The functional annotation of all peaks identified independently in BAP1, HCFC1 and FOXK2 ChIP-seq analyses revealed that, unlike cancer cell lines (Wang et al., 2018), BAP1 is primarily bound at genes promoters with a marginal association to putative distal regulatory sites in ESC (Figure 1C) in agreement with a recent report (Kolovos et al., 2020). While HCFC1 followed a similar behavior, FOXK2 localization showed a broader distribution that likely involved also enhancer occupancy (Figure 1C). Considering the high degree of overlap for FOXK2 with BAP1 bound promoters (Figure 1A), it’s likely that the activity of FOXK2 at distal elements involves other chromatin regulatory complexes unrelated to PR-DUB activity (Laget et al., 2010). We further explore the transcriptional state of target genes and found that, while RING1B targets were as expected largely inactive, BAP1 targets were instead actively transcribed (Figure 1D). This is in accordance with the lack of repressive histone post-translational modifications (PTMs) deposited by PRC1 and PRC2 (H3K27me3 and H2AK119ub1) and with the accumulation of H3K27ac at BAP1-unique promoters (Figure 1A). Together, these data illustrate that PR-DUB complex binding is largely independent of PRC1 and PRC2 complex (Figure 1E) and is primarily found at active promoters.

### BAP1 loss causes global increases in H2AK119ub1 and displacement of PRC1 from target loci

To investigate the effect of BAP1 loss of function on PRC1 complex activity and genomic occupancy, we generated stable BAP1 KO ESC by CRISPR Cas9 engineering. Western blot analysis of two independent BAP1 KO ESC lines showed the complete absence of BAP1 expression together with a global increase in H2AK119ub1 deposition (Figure S2A) as previously described (Campagne et al., 2019). RNA-seq analysis highlighted a preference in gene downregulation compared to activation (140 vs 29 for clone 1 and 174 vs 58 for clone 2; Figure 1F). Interestingly, such downregulation involved a very small number of BAP1 direct targets (Figure 1G), suggesting that BAP1 activity does not serve to counteract repressive signals like PcG activity at these sites and is dispensable to sustain direct target genes expression under homeostatic conditions. However, there is a higher number of PcG targets affected by BAP1 loss (Figure 1G) suggesting that BAP1 is more important for regulating these sites than its own target genes.

To further explore the molecular role of BAP1 and the consequences to its loss of function, we created a system where we can delineate the structural functions of BAP1 from its catalytic role. This involved the rescue of BAP1 loss of function by stable expression of either WT or catalytic BAP1 mutant (C91S) in KO ESCs (Figure S2B). Stable ESC lines using an empty pCAG-FLAG/HA were also generated in both parental and BAP1 KO ESC and served as further controls (Figure S2B). Western blot analysis in these models demonstrated that, while the expression of WT BAP1 restored normal H2AK119ub1 levels, the expression of the catalytic inactive C91S mutant did not. Importantly, this occurred in absence of major expression changes in all other PR-DUB subunits without affecting the expression levels of the core Polycomb components RING1B and SUZ12 as well as of the histone demethylase UTX that can specifically target H3K27me3 for PTM removal (Figure 2A and Figure S2A).

**Figure 2.**
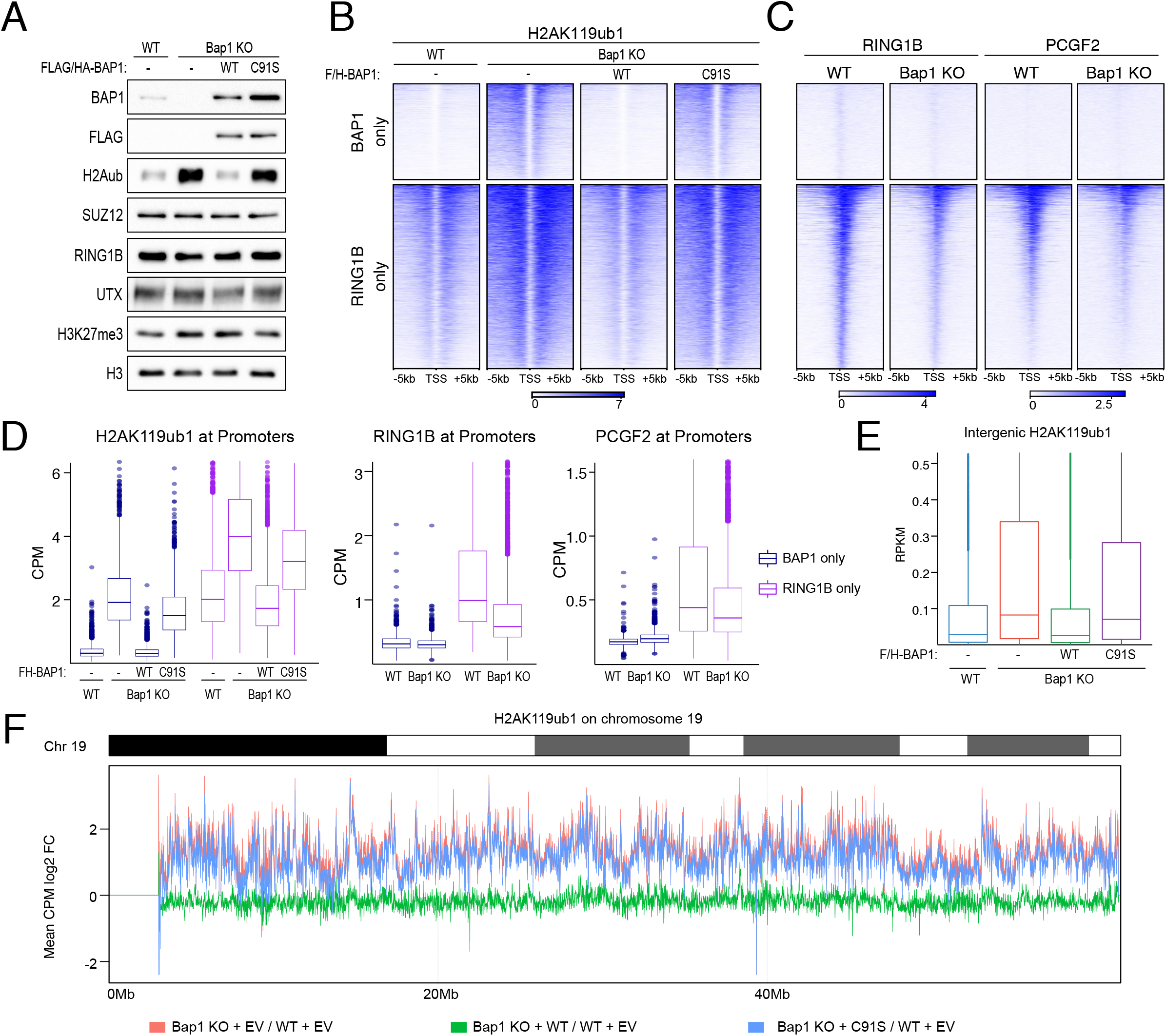
BAP1 loss causes global increases in H2AK119ub1 and displacement of PRC1 from target loci. A) Western blot analysis with the indicated antibodies on total protein extracts from the indicated rescue ESC cell lines (E14 WT + empty vector, Bap1 KO + empty vector, Bap1 KO + BAP1 WT, Bap1 KO + BAP1 C91S). B) Heatmaps representing normalized ChIP-seq intensity for H2AK119ub1 in the indicated cell lines +/− 5kb of TSS. Clusters are divided into BAP1 target genes only and RING1B only targets. C) Heatmaps representing ChIP-seq intensity for RING1B or PCGF2 in the indicated cell lines +/− 5kb of TSS. Clusters are divided into BAP1 target genes only and RING1B only targets. D) Boxplot of normalized intensity profiles for H2AK119ub1, RING1B and PCGF2 ChIP-seq in the indicated cell lines at BAP1 target genes only (blue) and RING1B only targets (purple). E) Boxplot representing H2AK119ub1 ChIP-seq RPKM levels in the indicated cell lines at intergenic sites. F) Representation of the Log2 Fold change CPM in H2AK119ub1 ChIP-seq signal in the indicated cell lines across chromosome 19 using 10kb windows.

To examine the effect of BAP1 KO and rescues on H2AK119ub1 deposition at genome wide-level, we performed quantitative spike-in ChIP-seq analyses (Figure 2B). This reveled that both BAP1-unique and RING1B-unique clusters presented extensive accumulation of H2AK119ub1 that was dependent on BAP1 catalytic activity (Figure 2B and D). The analysis of the H2AK119ub1 spreading, either 3’ or 5’ to the RING1B peak area, demonstrated that the gain in signal was greater outside the peak compared to inside (Figure S2C, D and E). This suggests that BAP1-dependent accumulation of H2AK119ub1 is caused by a spreading effect rather than a regulatory balance at PcG enriched sites. Consistently, the analysis of the binding for RING1B and PCGF2 revealed a reduction in occupancy for PRC1 core subunits at RING1B-unique targets that was dependent on BAP1 catalytic activity. Importantly, this occurred in the absence of any *de novo* gain in PRC1 association along the genome including at BAP1-unique target sites despite a clear accumulation in H2AK119ub1 levels (Figure 2C and 2D).

To further probe the genome wide effect on H2AK119ub1 levels, we quantified this PTM at intergenic sites demonstrating that H2AK119ub1 accumulated at all intergenic sites when BAP1 catalytic activity was lost (Figure 2E). This allows us to speculate that BAP1 activity is devoted to preventing aberrant H2AK119ub1 accumulation at the genome wide level. To further test this hypothesis at chromosome-wide level, we divided the entire mouse genome in 10kb windows and measured H2AK119ub1 levels within each bin. This revealed that H2AK119ub1 levels were significantly increased across the entire chromosomes (Figure 2F; chromosome 19 was shown as a representative example). Importantly, while re-expression of BAP1 restored normal H2AK119ub1 levels, the expression of its catalytic-inactive C91S mutant did not. Together, these analyses demonstrate that BAP1 maintains low level of extra-genic (extra-promoter) H2AK119ub1 deposition and that its loss seems to mobilize PRC1 from its physiological targets contributing to spurious non-targeted H2AK119ub1 deposition across the entire genome.

### PCGF3/5-PRC1 complexes are the major source of diffuse H2AK119ub1 deposition

While the knockout of RING1A/B has previously been shown to rescue BAP1 loss mediated transcriptional changes (Campagne et al., 2019), this approach remained a poor strategy to counteract pathological deficiencies of BAP1. RING1A/B activity is generally required for cell and adult tissue viability (Chiacchiera et al., 2016; Cohen et al., 2018; Pivetti et al., 2019; Voncken et al., 2003) since it touches the activity of multiple PRC1 sub-complexes (Fursova et al., 2019; Scelfo et al., 2019). Therefore, we decided to delve further into the precise relationship between the activity of BAP1 with respect to the distinct PRC1 sub-complexes in regulating H2AK119ub1 levels in order to uncover potential specificities.

Since BAP1 loss preferentially induced a genome-wide diffuse accumulation of H2AK119ub1 not clustered at specific loci, we reasoned that PCGF3 and PCGF5 containing complexes (PRC1.3/5) could play a critical role in such regulation. Diffuse deposition of H2AK119ub1 is reminiscent of the X-inactivation where PCGF3/5 were shown to be the only sub-complexes required for H2AK119ub1 deposition during XCI (Almeida et al., 2017). First, we generated a model in which the PRC1.3/5 activity is specifically lost together with BAP1 (PCGF3/5 KO +/− BAP1 KO). Second, we generated anther model in which PRC1.3/5 was the only PRC1 activity left in ESC (PCGF1/2/4/6 KO +/− BAP1 KO). Western blot analyses with these tools revealed that specific PCGF3/5 loss of function induced a general reduction in H2AK119ub1 levels (Figure 3A). Importantly, further loss of BAP1 was not sufficient to restore normal H2AK119ub1 levels in PCGF3/5 KO ESC (Figure 3A). Only a modest increase was observed when BAP1 was depleted suggesting that the majority, but not all, BAP1-opposed H2AK119ub1 is generated by PCGF3/5 containing complexes. In comparison, the PCGF1/2/4/6 knockout cells showed unaltered H2AK119ub1 levels, which were efficiently increased when BAP1 was lost similarly to their WT counterpart (Figure 3A). Together, these results strongly suggest that PCGF3/5 are primarily responsible for diffuse genome-wide accumulation of H2AK119ub1.

**Figure 3.**
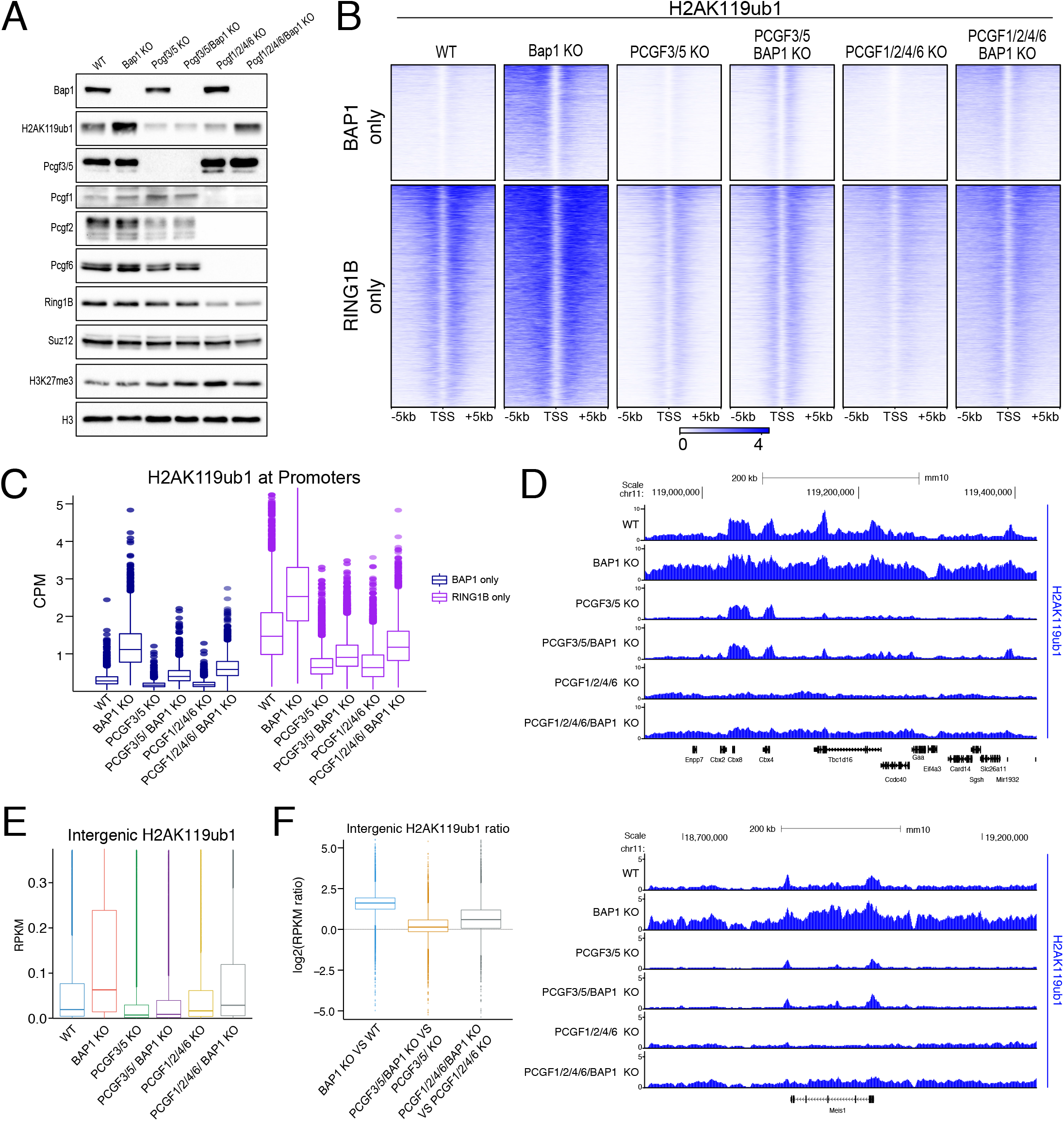
PCGF3/5-PRC1 complexes are the major source of diffuse H2AK119ub1 deposition. A) Western blot analysis with the indicated antibodies on total protein extracts from the WT, BAP1 KO, PCGF3/5 KO, PCGF3/5/BAP1 KO, PCGF1/2/4/6 KO and PCGF1/2/4/6/BAP1 KO ESCs. B) Heatmaps representing normalized ChIP-seq intensity for H2AK119ub1 in the indicated cell lines +/− 5kb of TSS. Clusters are divided into BAP1 target genes only and RING1B only targets. C) Boxplot of normalized intensity profiles for H2AK119ub1 ChIP-seq in the indicated cell lines at BAP1 target genes only (blue) and RING1B only targets (purple). D) UCSC genome browser snapshot of H2AK119ub1 ChIP-seq in the indicated cell lines. E) Boxplots representing H2AK119ub1 ChIP-seq RPKM levels in the indicated cell lines at intergenic sites. F) Boxplots showing the log2(Fold change RPKM) ratio for H2AK119ub1 in the indicated cell lines comparisons at intergenic regions.

To further extend these observations, we performed ChIP-seq analyses for H2AK119ub1 using these lines. This revealed different patterns of regulation for H2AK119ub1 deposition at RING1B target promoters (Figure 3B and C). Quantification of H2AK119ub1 levels at promoters were apparently decreased in both PCGF3/5 and PCGF1/2/4/6 KO models (in presence of BAP1). However, while loss of PRC1.3/5 affected spreading but not the normal bimodal accumulation of H2AK119ub1 at PcG target TSS; in the presence of PRC1.3/5 activity only (PCGF1/2/4/6 KO) such accumulation was lost and converted into diffuse H2AK119ub1 deposition (Figure 3B and D). The removal of BAP1 did not lead to any evident increase in diffuse H2AK119ub1 deposition in the absence of PCGF3/5. Consistently, when only PRC1.3/5 sub-complexes were active in ESC, BAP1 removal specifically increased the diffuse deposition of H2AK119ub1 (Figure 3D). Indeed, the quantification of H2AK119ub1 levels at all intergenic regions followed this trend (Figure 3E and 3F), demonstrating that PRC1.3/5 activity is primarily responsible for the large-scale intergenic changes in H2AK119ub1 upon *Bap1* deletion in a fashion resembling the spread of H2AK119ub1 on the inactive X-chromosome (Almeida et al., 2017).

### BAP1 catalytic activity maintains stable PRC2 target association and spatial distribution of H3K27me3

There is conflicting evidence on the role played by PRC2 activity with respect to BAP1 tumor suppressive function (Abdel-Wahab et al., 2012; Lafave et al., 2015; Schoumacher et al., 2016). We and others have recently shown that H2AK119ub1 provides an important contribution to the stable binding of PRC2 to its target genes (Blackledge et al., 2014; Healy et al., 2019; Tamburri et al., 2020) as well as for its catalytic activity (Kalb et al., 2014; Kasinath et al., 2020). Thus, we performed ChIP-seq analyses for H3K27me3 and SUZ12 in our rescue system and found that H3K27me3 deposition followed H2AK119ub1 only to some extent. At BAP1-unique targets, loss of BAP1 or its catalytic activity caused a gain of H3K27me3 deposition 5’ to the TSS that was independent of *de novo* SUZ12 binding (Figure 4A and B). The lack of 3’ invasion of H3K27me3 follows H2AK119ub1 deposition and is likely a consequence of an antagonistic relationship with RNA polymerase activity which maintains active transcription of these genes (Figure 1C; Beltran et al., 2016; Riising et al., 2014; Zhang et al., 2019). More interestingly, at RING1B target genes, we observed that both H3K27me3 deposition and SUZ12 binding was reduced in absence of BAP1 (Figure 4A and 3B) despite the extensive gains in H2AK119ub1. Importantly, such displacement is dependent on BAP1 catalytic activity as it was not rescued by C91S expression (Figure 4A and 4B). This result is consistent with the requirement of ASXL1 for correct PRC2 binding and activity at the HOX locus genes in leukemia cells (Abdel-Wahab et al., 2012).

**Figure 4.**
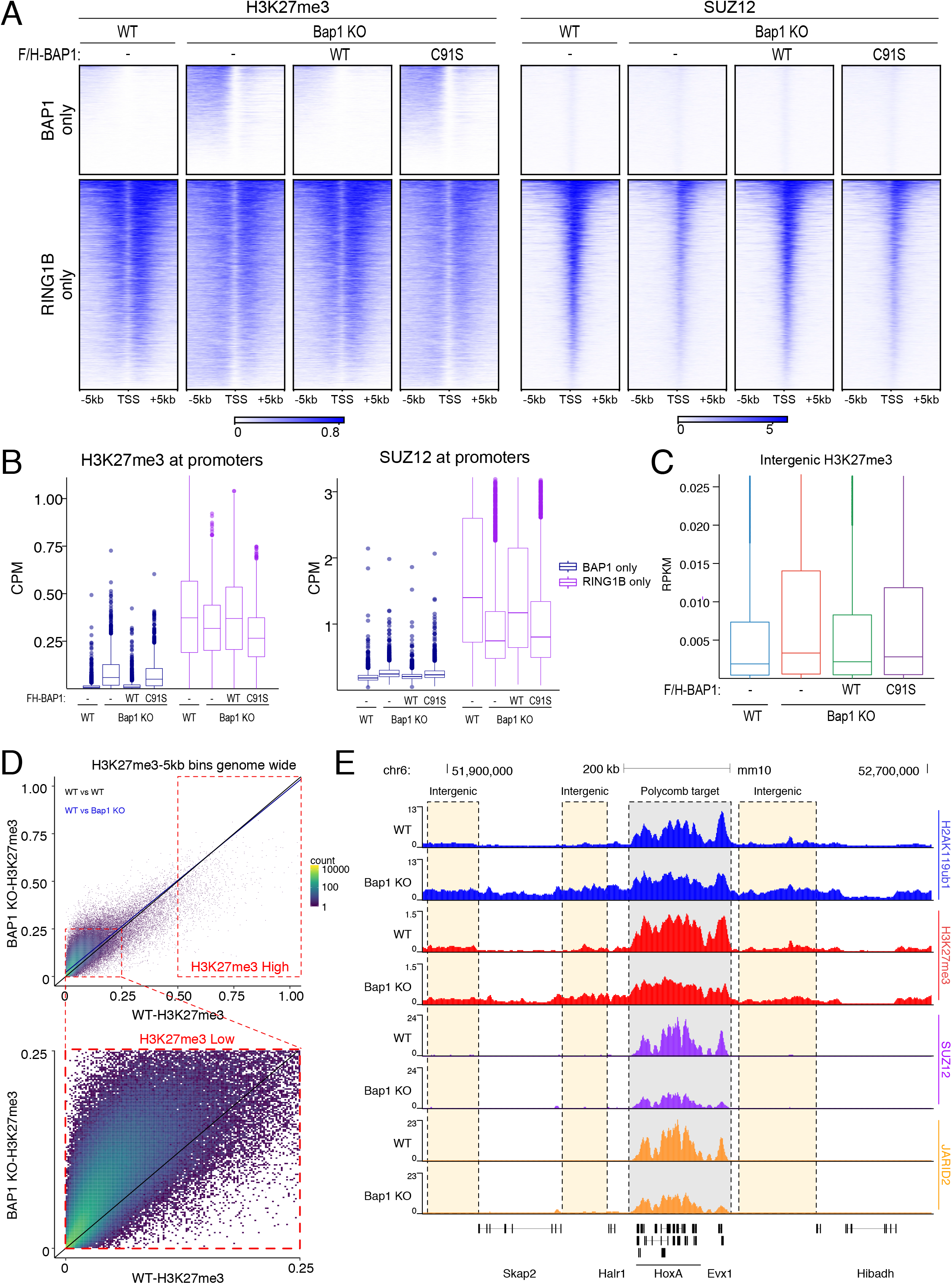
BAP1 catalytic activity maintains stable PRC2 target association and spatial distribution of H3K27me3. A) Heatmaps representing normalized ChIP-seq intensity for H3K27me3 (left), or ChIP-seq intensity for SUZ12 (right) in the indicated cell lines +/− 5kb of TSS. Clusters are divided into BAP1 target genes only and RING1B only targets. B) Boxplot of normalized intensity profiles for H3K27me3 and SUZ12 ChIP-seq in the indicated cell lines at BAP1 target genes only (blue) and RING1B only targets (purple). C) Boxplots representing H3K27me3 ChIP-seq RPKM levels in the indicated cell lines at intergenic sites. D) Genome-wide XY scatterplot of normalized H3K27me3 ChIP-seq intensities at a resolution of 5kb in Bap1 KO compared to WT ESCs. Each point represents one 5kb window. E) UCSC genome browser snapshot of indicated ChIP-seq tracks at the HOXA locus in WT and Bap1 KO ESCs. Highlighted regions show Polycomb targets (grey) or intergenic regions (yellow) showing the trend of redistributed H3K27me3 from Polycomb targets to intergenic sites in Bap1 KO ESCs.

Similar to BAP1-unique targets, quantifications at all intergenic sites showed that H3K27me3 underwent accumulation with a similar trend to H2AK119ub1 in absence of BAP1 activity. Further analysis of the entire genome divided into 5kb windows confirmed this result, demonstrating that H3K27me3 accumulated exclusively at genomic regions with low levels for this PTM (Figure 4D). In contrast, the few regions that retained high levels of H3K27me3 (likely PcG target loci) reduced this modification (Figure 4D) in agreement with displacement of PRC2 from its target sites. Overall, this suggests that intergenic gain in H2AK119ub1 titrates PRC2 away from its normal target promoters, decreasing promoter H3K27me3 concentration and increasing its indiscriminate intergenic levels (Figure 4E).

### AEBP2 and JARID2 are required for intergenic H3K27me3 titration

The PRC2.2 sub-complex specific subunits JARID2 and AEBP2 have been widely reported to have an affinity for H2AK119ub1 (Cooper et al., 2016; Kalb et al., 2014; Kasinath et al., 2020). This affinity is essential for PRC2 recruitment to the inactive X-chromosome (Cooper et al., 2016) and removal of H2AK119ub1 results in a preferential loss of PRC2.2 binding compared to PRC2.1 (Blackledge et al., 2020; Tamburri et al., 2020). Since the affinity of PRC2.2 for H2AK119ub1 could play a role in titrating away PRC2 from its target promoters, we knocked out AEBP2 and JARID2 in the presence (A+J) or absence of BAP1 (A+J+B; Figure 5A). As expected, H2AK119ub1 levels were increased to a similar extent in the presence or absence of AEBP2 and JARID2, demonstrating that the gain in PRC1 activity occurs upstream of PRC2.2 (Figure 5A). We performed ChIP-seq analysis for both H2AK119ub1 and H3K27me3 and observed similar changes in BAP1 and AEBP2/JARID2/BAP1 KO cells at RING1B target genes (Figure 5B and 5C). This showed that AEBP2 and JARID2 removal did not rescue the reduction in H3K27me3 levels at PcG targets. However, analysis of intergenic regions showed that loss of AEBP2 and JARID2 uncouples the gain in non-genic H3K27me3 deposition from H2AK119ub1 accumulation (Figure 5D and S3A). While the fold change in H2AK119ub1 accumulation in non-genic regions remained similar in the presence or absence of AEBP2 and JARID2; BAP1 loss did not lead to any additive accumulation in H3K27me3 in the absence of AEBP2 and JARID2 (Figure 5D). Please note that there is already a general gain in intergenic H3K27me3 levels in AEBP2/JARID2 KO cells (Figure S3B and S3C). This is in line with previous reports (Højfeldt et al., 2019) and is the results of a reduced tethering of core PRC2 to target promoters, which is independent of intergenic H2AK119ub1 accumulation.

**Figure 5.**
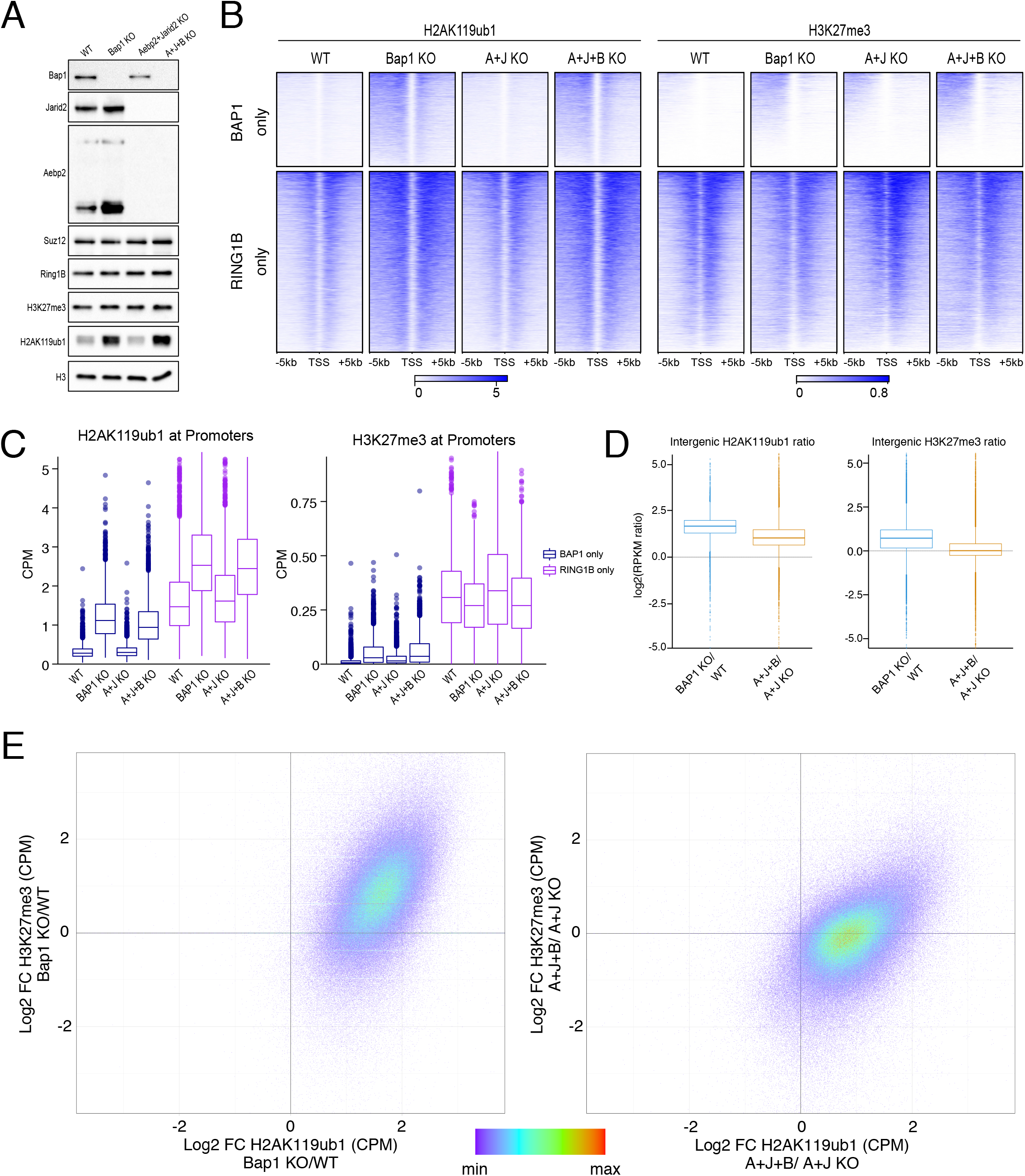
AEBP2 and JARID2 are required for intergenic H3K27me3 titration. A) Western blot analysis with the indicated antibodies on total protein extracts from the WT, BAP1 KO, Aebp2+Jarid2 (A+J) KO, Aebp2+Jarid2+Bap1 (A+J+B) KO ESCs. B) Heatmaps representing normalized ChIP-seq intensity for H2AK119ub1 (left) or H3K27me3 (right) in the indicated cell lines +/− 5kb of TSS. Clusters are divided into BAP1 target genes only and RING1B only targets. C) Boxplot of normalized intensity profiles for H2AK119ub1 and H3K27me3 ChIP-seq in the indicated cell lines at BAP1 target genes only (blue) and RING1B only targets (purple). D) Boxplots showing the log2(Fold change RPKM) ratio for either H2AK119ub1 (left) or H3K27me3 (right) in the indicated cell lines comparisons at intergenic regions. E) Genome wide comparison of ChIP-seq signal using 5kb windows. Log2 fold change H2AK119ub1 ChIP-seq for the indicated cell lines (X-axis) plotted against Log2 fold change of H3K27me3 ChIP-seq (Y-axis). Each dot represents one 5kb window.

The uncoupling of intergenic H3K27me3 gain from H2AK119ub1 spreading can be better visualized through scatter plots of the entire genome in 5kb windows, where Log2 fold changes in H2AK119ub1 were plotted against H3K27me3 (Figure 5E). Importantly, this analysis demonstrated a high concordance in the genomic regions that gained both H2AK119ub1 and H3K27me3 (Figure 5E, left panel). When the same analysis was performed in absence of AEBP2/JARID2, H2AK119ub1 accumulation was clearly uncoupled from H3K27me3 (Figure 5E, right panel). This supports our hypothesis that JARID2 and AEBP2 are required for the titration of PRC2 activities away from their target sites in a BAP1 null context.

### Diffuse H2AK119ub1 accumulation causes global chromatin compaction

To this point we have revealed the mechanism of reshaping the PRC1 and PRC2 mediated landscape in BAP1 KO ESC. However, we wanted to investigate how the spreading of Polycomb modifications and the titration of PRC2.2 from its target sites may affect chromatin structure to further understand the mechanisms perturbed in patients with BAP1 inactivating mutations. Since the activity of the PRC1.3/5 complexes has been linked to large scale chromatin compaction through XCI (Almeida et al., 2017), we wondered whether diffuse accumulation of H2AK119ub1 coupled to H3K27me3 deposition could lead to changes in 3D chromatin organization and compaction. For this, we performed *in situ* Hi-C analysis in WT and *Bap1* KO ESC and found that loss of BAP1 induces a general gain in contacts across the entire genome as exhibited in the representative contact matrices for chromosome 11 (Figure 6A). Upon splitting these contact changes into quartiles based on the Log2 fold change versus WT, all four quartiles have an average enrichment for BAP1 KO (Figure 6B). This allows us to conclude that over three-quarters of the genome acquired a more compact configuration in the absence of BAP1. Correlation of changes in histone modifications within these quartiles in BAP1 KO vs WT ESCs showed that the largest gain in H2AK119ub1 and H3K27me3 was exhibited at the mid-high quartiles, coinciding with the largest loss in H3K4me3 (Figure 6C). This hints at a model where the broad gains in Polycomb modifications coincide with a reduction in active histone modifications. Overlaying Hi-C fold change matrices with ChIP-seq profiles highlighted the spreading of H2AK119ub1 and H3K27me3 relative to compaction and some local reductions in H3K4me3 deposition (Figure 6D). Importantly, loss of BAP1 only affected the general compaction state of chromatin without altering the status and boundaries of topology associated domains (TADs).

**Figure 6.**
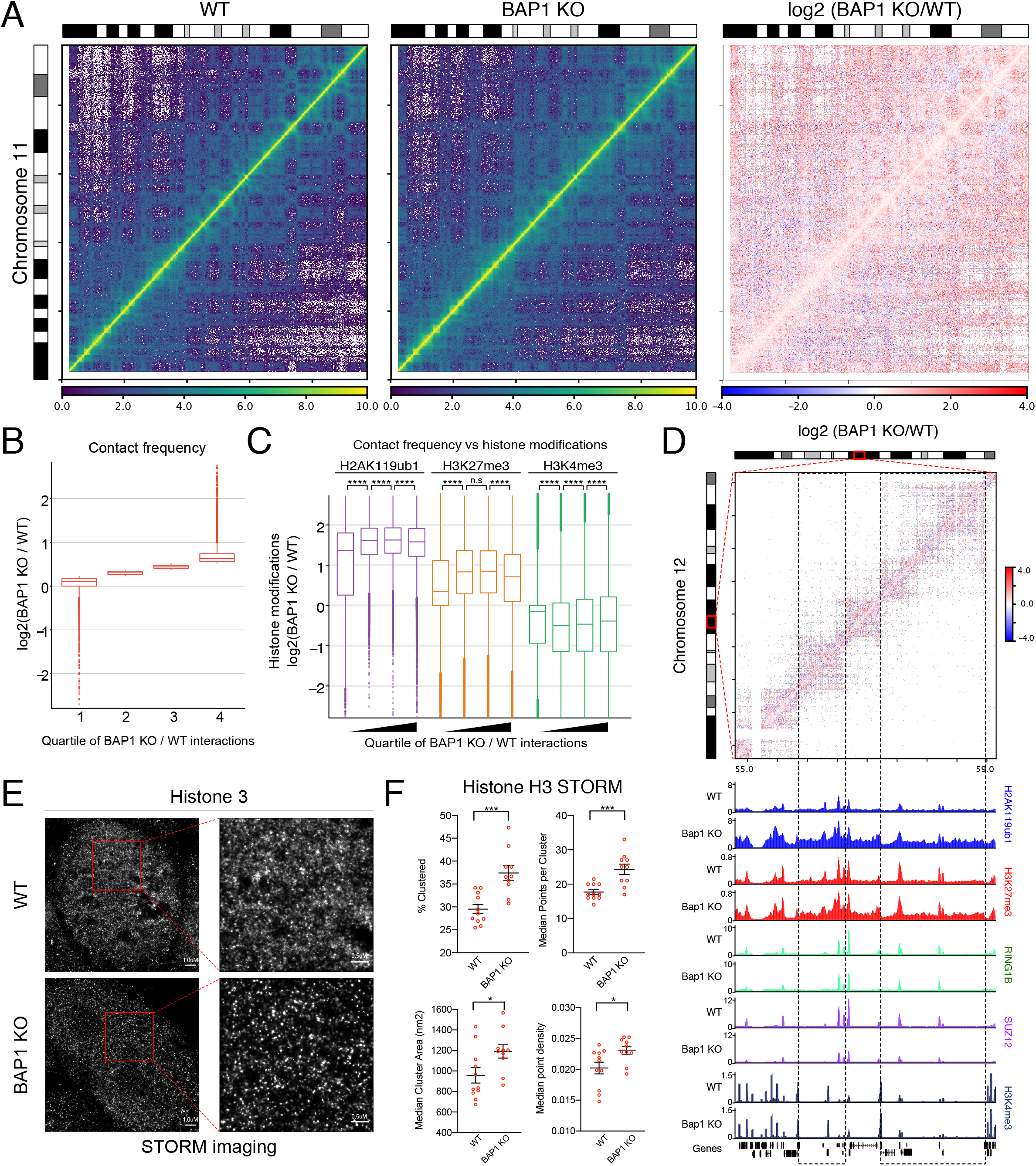
Diffuse H2AK119ub1 accumulation causes global chromatin compaction. A) HiC contact matrix of the entire chromosome 11 in WT, BAP1 KO and log2 (Fold change BAP1 KO/WT) at 250kbp resolution. B) Boxplot of contact frequency of log2(BAP1 KO/WT) ratios divided into quartiles. C) Boxplot of the log2(BAP1 KO/WT) ratio of the indicated histone modifications within the quartiles defined in B. Wilcoxon test was used to ascertain significance. D) Top. Log2 (fold change BAP1 KO/WT) HiC contact matrix of the indicated region of chromosome 12 (54.8-59.15Mb) at 10kb resolution. Bottom. UCSC genome browser snapshot of indicated ChIP-seq tracks at the same region of chromosome 12. This highlights the trend observed in C of spreading of H2AK119ub1 and H3K27me3 within TADs, with focal reductions in H3K4me3 at certain peaks. E) Representative Stochastic Optical Reconstruction Microscopy images of WT or BAP1 KO ESC stained with Histone 3. F) STORM quantifications of Histone H3 features using multiple regions of interest for WT (n=11) and BAP1 KO (n=10) ESCs.

Quantitative analysis by mass spectrometry of chromatin associated proteins revealed an enrichment for repressive and heterochromatic factors upon BAP1 loss coupled with a displacement of factors associated with active transcription and replication. This included accumulation of HIST1H1E, HIST1H1D, DNMT3A/B/L and CBX3 as well as loss of POLR1A/B, POLR2H, ERCC2/3, SMARCD2, TOP1 and MCM7 chromatin association (Figure S4A). Interestingly, Histone H1 has recently been shown to promote spreading of H2AK119ub1 through compacting nucleosomes (Zhao et al., 2020), supporting a model of chromatin compaction that facilitates broad H2AK119ub1 spreading. Together, this further supports the possibility that loss of BAP1 creates a more compact chromatin environment.

*In vivo* nucleosomes are organized in cluster (clutches) of different sizes interspersed by nucleosome-free DNA. The size of clutches can vary from cell type to cell type with pluripotent cells showing small and diffuse nucleosome clusters and more committed cell types a closer chromatin conformation characterized by larger and more dense nucleosomal clusters (Ricci et al., 2015). We hypothesize that changes in chromatin compaction should reflect in a different density of clutches distribution. Therefore, we performed Histone H3 staining followed by Stochastic Optical Reconstruction Microscopy (STORM) imaging to identify and quantify nucleosome clutches in BAP1 WT vs. KO ESC. Consistent with a tighter chromatin conformation revealed by Hi-C, representative images of STORM reconstruction highlighted that the chromatin of *Bap1* KO ESC displayed brighter and less diffused nucleosome clusters (Figure 6E). Quantification of Histone H3 molecules (point) localisation and grouping into clusters (Figure S4B) further demonstrated that histone H3 becomes more densely packed when BAP1 activity is lost (Figure 6F). This included an increased percentage of clustered molecules (points) forming an average of larger clusters compared to WT ESC (Figure 6F). Together, these results demonstrate that BAP1 preserves low levels of H2AK119ub1 across the genome to counteract a general compaction of chromatin.

We propose a model (Figure 7A) where the primary role of BAP1 is in removing non-specific PCGF3/5 dependent H2AK119ub1 throughout the genome. The absence of BAP1 leads to accumulation of H2AK119ub1 intergenically which titrates PRC2 away from its target genes through JARID2 and AEBP2. This depletion of Polycomb complexes from their target sites allows aberrant activation of transcription, while simultaneously causing repressive chromatin compaction elsewhere. This indirect regulation of transcription and chromatin conformation may help explain the tumor suppressive functions of BAP1 that are lost in BAP1 mutant cancers.

**Figure 7.**
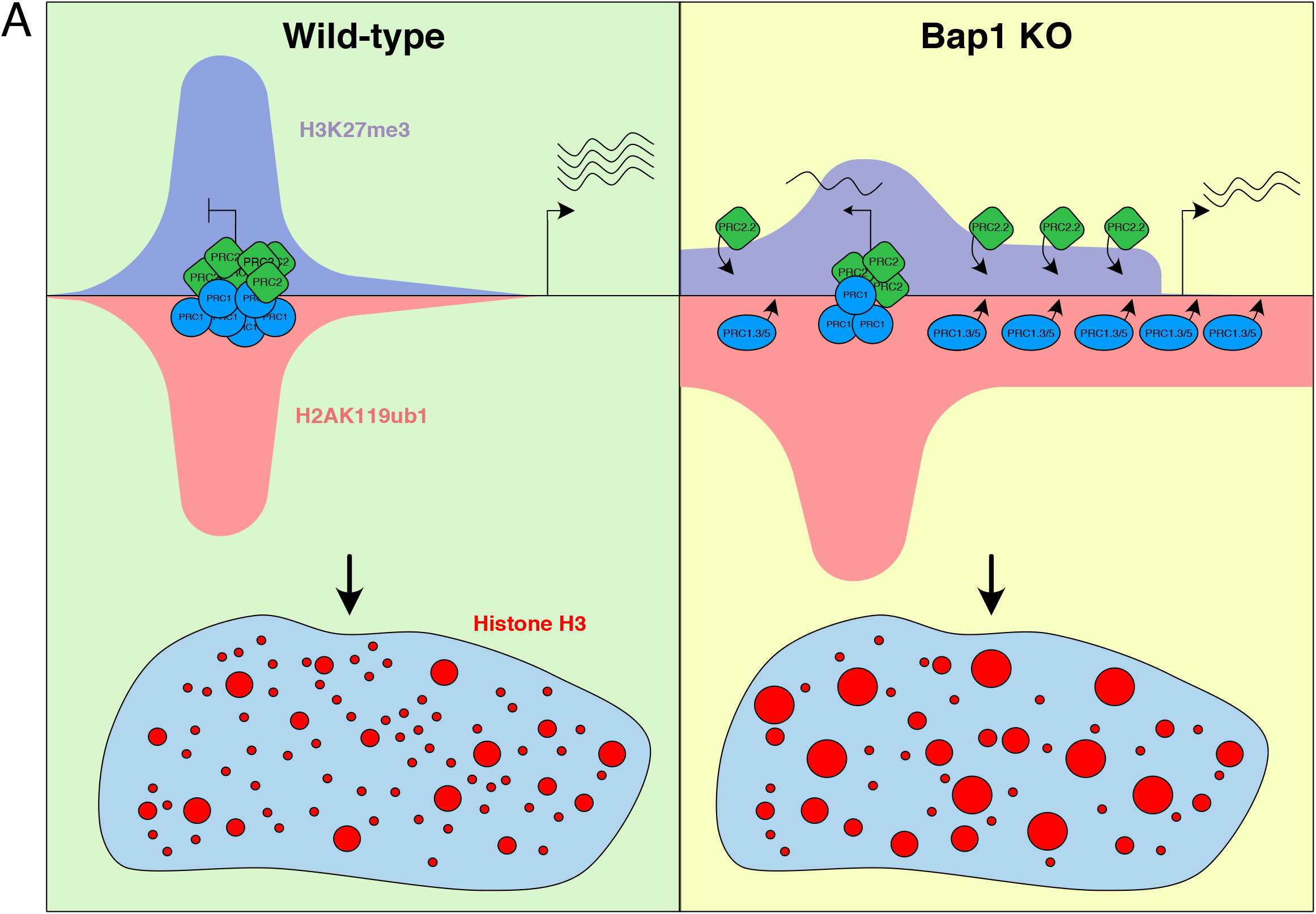
Mode of action of BAP1. A) Model of the dual role of BAP1 mode of action on transcription. BAP1 is essential for the spatial maintenance of H2AK119ub1 and H3K27me3. Spurious redistribution of these in the absence of BAP1, directed by the PCGF3/5-PRC1 and PRC2.2 complexes, promotes chromatin compaction and a general repression of transcription (Trithorax phenotype) while simultaneously allowing derepression of selected Polycomb target genes (Polycomb phenotype).

## Discussion

Here we have provided evidence that uncover the mode of action of BAP1 in the field of chromatin biology which can be highly relevant to disease, proposing a mechanism of how loss-of function BAP1 mutants may disrupt transcriptional programs in tumorigenesis. Activation of Polycomb target genes in the absence of BAP1 mimics PRC2 loss-of-function mutations and events, which are a common feature of many cancers such as T-ALL, MPNST, Ependymoma, DIPG and others (Conway et al., 2015; Jain et al., 2019; Lee et al., 2014; Ntziachristos et al., 2012; Piunti et al., 2017). This loss of Polycomb silencing is precisely what is observed in ASXL1 mutant myeloid leukemias also, as loss of ASXL1 removes PRC2 from its promoters, allowing activation of oncogenic activities (Abdel-Wahab et al., 2012). Titration of PRC1 and PRC2 from their canonical sites is an emerging theme in the chromatin field as it has now been shown that deletion of BAF complex subunits paradoxically reduces Polycomb complex occupancy and silencing through the titration of PRC1/2 from their target sites (Weber et al., 2020). As the subunits of the BAF complex, like PR-DUB, are also frequently mutated in a number of cancers (Bracken et al., 2019), these new data suggest alternate mechanisms outside of PRC2 subunit mutation which can cause pathogenic PRC2 loss-of-function.

Curiously, the role of BAP1 in preventing intergenic spreading of H3K27me3 and chromatin compaction is reminiscent of EZH2 gain-of-function cancers such as non-Hodgkin lymphoma (Morin et al., 2011) in which there is a genome-wide gain in H3K27me3 levels that correlates with local increases in compaction and transcriptional inactivation of affected TADs (Donaldson-Collier et al., 2019). This paradigm of intergenic spreading of H3K27me3 is found in several other cancer types such as osteosarcomas featuring H3.3G34 mutations where the diffuse H3K27me3 represses distal regulatory elements such as enhancers (Jain et al., 2020).

The relevance of these findings is not limited to cancer, as neurodevelopmental disorders featuring mutations in the ASXL subunits of PR-DUB may also be affected through the same mechanisms (Bainbridge et al., 2013; Hoischen et al., 2011). Intriguingly, one of the few PRC1 subunits mutated in neurological and intellectual disability disorders is AUTS2, which is part of the PCGF3/5 complexes (Gao et al., 2014; Kalscheuer et al., 2007; Si et al., 2016). The antagonism we have demonstrated between BAP1 and PCGF3/5 complexes might suggest that a careful balance in the levels of intergenic or broad domains of H2AK119ub1 is essential to maintain transcriptional homeostasis during neurodevelopment.

It remains to be defined whether the contribution of BAP1 to tumor suppression is through the maintenance of Polycomb repression or through the inhibition of repressive chromatin compaction. None-the-less our thorough genetic analysis of the mechanisms behind the intergenic spreading of H2AK119ub1 provides a number of potential synthetic lethalities that are worthy of further investigation in a cancer context. Foremost among these are the PCGF3/5 subunits, and potentially their specific complex members (Gao et al., 2012), along with AEBP2 and JARID2.

While it is clear that Polycomb complex mediated histone modifications are actively involved in transcriptional repression (Blackledge et al., 2020; Pengelly et al., 2013; Tamburri et al., 2020), it is not yet fully understood how exactly H2AK119ub1 participates in this. Although H2AK119ub1 does promote recruitment of PRC2.2 complexes, loss of PRC2 in a static system such as ESCs has very little effect on transcription, while loss of H2AK119ub1 has huge transcriptional effects, causing widespread activation of Polycomb targets (Blackledge et al., 2020; Lavarone et al., 2019; Tamburri et al., 2020). Therefore, there is some underappreciated role of H2AK119ub1 in blocking transcription, perhaps through its antagonistic relationship with RNA PolII initiation (Dobrinić et al., 2020). It is possible that this transcriptional silencing function of H2AK119ub1 causes widespread repressive effects in BAP1 mutant cancers which causes the pathogenic effects on transcription. The role of PRC1 in regulating the 3D epigenome has been investigated primarily through non-catalytic roles of canonical PRC1 and the compaction related functions of the PHC and CBX subunits (Isono et al., 2013; Kundu et al., 2017; Plys et al., 2019; Tatavosian et al., 2019). Whether H2AK119ub1 actually plays a role in this process has been unclear due to the use of hypomorphic mutants of RING1B to date (Boyle et al., 2020; Kundu et al., 2017). The essential role of PCGF3/5 in X-chromosome inactivation, and the requirement of JARID2 for XCI both suggest that H2AK119ub1 is involved in this type of compaction (Almeida et al., 2017; Cooper et al., 2016). However, whether that role extends to somatic chromosomes and local 3D structures such as TADs and Polycomb bodies is not yet understood. Our data suggest that diffuse intergenic levels of H2K119ub1 can stimulate local compaction of chromatin in a repressive fashion, in similar way to XCI.

Finally, the dual function we have described for BAP1 in both the activation and repression of Polycomb target genes helps resolve a puzzling issue in developmental biology. The apparent Polycomb and Trithorax *in vivo* phenotype in Calypso and ASXL mutants in Drosophila and mice (Baskind et al., 2009; Fisher et al., 2010; Scheuermann et al., 2010) have been confounding mechanistically until now. It is clear that in the developmental context that a fine balance between H2AK119ub1 levels is essential to maintain normal gene expression programs.

## Methods

### Cell lines and cell culture

mESCs were grown on 0.1% gelatin-coated dishes in 2i/LIF-containing GMEM medium (Euroclone) supplemented with 20% fetal calf serum (Euroclone), 2 mM glutamine (Gibco), 100 U/ml penicillin, 0.1 mg/ml streptomycin (Gibco), 0.1 mM non-essential amino acids (Gibco), 1 mM sodium pyruvate (Gibco), 50 μM ß-mercaptoethanol phosphate buffered saline (PBS; Gibco), 1000 U/ml leukemia inhibitory factor (LIF; produced in-house), and GSK3β and MEK 1/2 inhibitors (ABCR GmbH) to a final concentration of 3 μM and 1 μM, respectively. Indicated cells were treated for 48 hours with 500nM dTAG-13 (TOCRIS), or DMSO as vehicle, in order to degrade dTAG-BAP1.

To generate stable KO cell lines, 10ug pX458 2.0 plasmid pairs (Addgene) encoding Cas9 and sgRNAs were transfected using Lipofectamine 2000 (Invitrogen), according to manufacturer’s instruction. Sorting of GFP positive cells was carried out 48 hours after transfection and 1000 cells were seeded onto a 15-cm dish. Clones were isolated 10-14 days later and grown further before screening for knockout by Western blot.

For rescue clone generation, mESCs were transfected with 10ug pCAG vectors encoding 2xFlag-HA-tagged BAP1 wild-type or BAP1 C91S using Lipofectamine 2000 (ThermoFisher Scientific), according to manufacturer’s instructions. 24 hours post-transfection puromycin selection (1μg/ml) was added for a further 24 hours. Cells were then split to clonal density (~1:40) onto a 15cm plate. Clones were isolated 10-14 days later and grown further before screening for rescue allele expression by Western blot.

### gRNA sequences

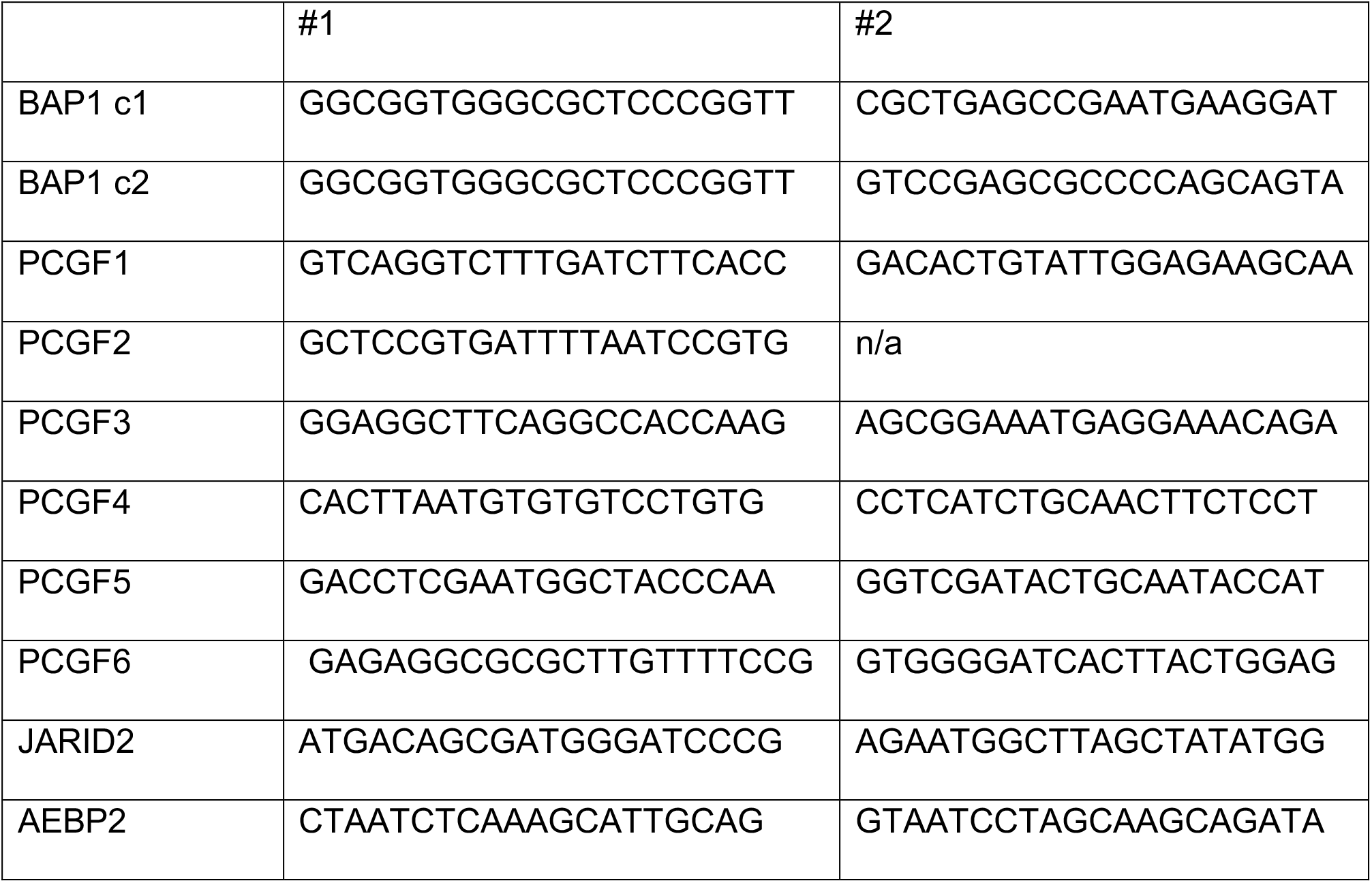

### Western Blot

For western blot analysis on total protein lysates, mESCs were lysed and sonicated in ice-cold S300 buffer (20 mM Tris-HCl pH 8.0, 300 mM NaCl, 10% glycerol, 0.2% NP40) and supplemented with protease inhibitors (Roche). Precipitates were removed by centrifugation. Lysates were then resuspended in Laemmli sample buffer and boiled for 5 minutes. Protein lysates were separated on SDS-PAGE gels and transferred to nitrocellulose membranes. After probing with the suitable primary and secondary antibodies, chemiluminescence signals were captured with the ChemiDoc Imaging System (Bio-Rad).

### Antibodies

Western blot and ChIP analyses were performed with: anti-BAP1 (D7W70; Cell Signaling), anti-Suz12 (D39F6; Cell Signaling Technology), anti-FLAG (F3165; Sigma-Aldrich), anti-H3K27me2 (9728; Cell Signaling Technology), anti-H3K27me3 (9733; Cell Signaling Technology), anti-H2AK119ub1 (8240; Cell Signaling Technology), anti-H3 (ab1791; Abcam), anti-H2A (07-146; Sigma-Aldrich), anti-RING1B (homemade (Chiacchiera et al., 2016)), anti-PCGF1, anti-PCGF2, and anti-PCGF6 (Pasini’s lab, homemade (Scelfo et al., 2019)), anti-JARID2 (13594, Cell Signaling Technology), anti-AEBP2 (14129; Cell signaling Technology), anti-HA (C29F4, Cell Signaling Technology), anti-ASXL2 (A302-037A, Bethyl), anti-ASXL1 (D1B6V, Cell Signaling Technology) anti-KDM1B/LSD2 (E1R60, Cell Signaling Technology), anti-OGT (SC-32921, Santa-Cruz), anti-HCFC1 (A301-400A, Bethyl), anti-FOXK2 (A301-730A, Bethyl), anti-MBD6 (SAB1305225-40TST, Sigma-Aldrich), anti-H3K4me1 (AB8895, Abcam), anti-H3K27ac (AB4729, Abcam) anti-PCGF3/5 (AB201510, Abcam), anti-UTX (Helin lab, homemade (Agger et al., 2007)).

### Chromatin immunoprecipitation (ChIP)

ChIP experiments were performed according to standard protocols as described previously (Ferrari et al., 2014). For all ChIPs, except HA, 1% formaldehyde cross-linked chromatin was sheared to 500–1000 bp fragments by sonication. For HA ChIPs cells were fixed in 2.5mM EGS for 50 minutes before the addition of formaldehyde to 1% for a further 10 minutes, before proceeding to sonication. Chromatin was then incubated overnight in IP buffer (33 mM Tris-HCl pH 8, 100 mM NaCl, 5 mM EDTA, 0.2% NaN_3_, 0.33% SDS, 1.66% Triton X-100) at 4°C with the indicated antibodies (5 μg antibodies/ 500 μg chromatin). For histone modifications ChIPs, 250 μg of chromatin supplemented with 5% spike-in of S2 Drosophila chromatin (prepared in the same manner) and 3 μg of antibodies was used. The next day, chromatin lysates were incubated for 3 hours with protein-G sepharose beads (GE Healthcare). Beads were washed 3× with low-salt buffer (150 mM NaCl, 20 mM Tris-HCl pH 8, 2 mM EDTA, 0.1% SDS, 1 % Triton X-100) and 1× with high-salt buffer (500 mM NaCl, 20 mM Tris-HCl pH 8, 2 mM EDTA, 0.1% SDS, 1% Triton X-100), and then re-suspended in de-crosslinking solution (0.1 M NaHCO3, 1% SDS). DNA was purified with QIAquick PCR purification kit (Qiagen) according to manufacturer’s instructions. DNA libraries were prepared with 2–10 ng of DNA using an in-house protocol (Blecher-Gonen et al., 2013) by the IEO genomic facility and sequenced on an Illumina HiSeq 2000.

### ChIP-seq data analysis

Paired-end DNA reads were processed through fastp to trim adapters and to remove low quality nucleotides at read ends (Chen et al., 2018). Quality-filtered DNA reads were aligned to the mouse reference genome mm10, or mm10 and fly reference genome (dm6) for histone ChIP-Rx using Bowtie v1.2.2 retaining only uniquely aligned reads (-m 1) and using the parameters -I 10, -X 1000 (Langmead et al., 2009). Reads mapped to both mm10 and dm6 were discarded. Peaks were identified using MACS2 v2.1.1 in narrow mode with parameters –format BAMPE –keep-dup all -m 3 30 and p-value 1e-10 (Zhang et al., 2008). Peaks were annotated using the R package ChIPpeakAnno v3.15 using for each peak the 5 kbp region around the center of the peak (Zhu et al., 2010).

The intensity of histone modifications or transcription factor binding was represented through boxplots or heatmaps, both generated from BigWig files that were obtained using the function bamCompare from deepTools 3.1 (Ramírez et al., 2016) with parameters – binSize 50 –extendReads. The –scaleFactors parameter of bamCompare was set to (1/total mapped reads)*1,000,000 to normalize for the differences in sample library size. Samples obtained with the ChIP-Rx method, were normalized using (1/dm mapped reads)*1,000,000 as –scaleFactors parameter (Orlando et al., 2014). ChIP-Rx samples were also normalized by the mm10/dm6 mapped reads of the input samples. A single genomic DNA input was sequenced for each cell line and used for the normalization of the respective ChIP-Rx samples. The preparation of the heatmaps required the generation of a data matrix through computeMatrix with parameters –referencePoint TSS/center -a 5000 -b 5000 (Ramírez et al., 2016). The data matrix was converted into heatmap by plotHeatmap (Ramírez et al., 2016). Boxplots were prepared using multiBigwigSummary in BED-file mode and using as bed file the regions corresponding to promoters (TSS +/− 2.5 kbp) or distal genomic regions such as enhancers (Ramírez et al., 2016).

Density plots were prepared using as input the intensity of histone modifications on 5 kbp windows at genome-wide level. When the log2 ratio between two conditions was reported, “Inf” and “-Inf” values were equaled to the maximum and minimum finite values, respectively. Venn diagram were prepared by the R package VennDiagram v1.6.20 using as input the name of the genes targeted by a specific protein or histone modification (Chen and Boutros, 2011).

### RNA-seq and analysis

RNA-seq was performed following SMART-seq2 protocol (Picelli et al., 2014) with minor modifications. Briefly, poly-A containing mRNA molecules obtained from 1 μg of total RNA were copied into first-strand cDNA by reverse transcription and template-switching using oligo (dT) primers and an LNA-containing template-switching oligo (TSO). Resulting cDNA was pre-amplified with KAPA HotStart Taq enzyme (Kapa Biosystems) and then purified with Ampure beads (Agencourt AMPure XP-Beckman Coulter). One nanogram of pre-amplified cDNA was tagmented with in-house produced Tn5 transposase and further amplified with KAPA HotStart Taq enzyme. After purification with Ampure beads, the quality of the obtained library was assessed by Bioanalyzer (High Sensitivity DNA kit, Agilent Technologies), prior to sequencing.

RNA reads were aligned to the mouse reference genome mm10 using STAR v2.7 without allowing for multimapping. PCR duplicates were removed using samblaster (Faust and Hall, 2014). Mapped reads were assigned to genes using featureCounts (Liao et al., 2014) with unstranded read counting on exons and using the gene name as attribute type in the annotation. Genes were annotated as in Gencode M21 (GRCm38) downloaded from https://www.genecodegenes.org/mouse/. Differentially expressed genes were identified using the R package DESeq2 v1.24 using default parameters (Love et al., 2014). The fold change of lowly expressed genes was corrected using the lfcShrink function of the apeglm R package with the type option set to apeglm v1.6 (Zhu et al., 2019). The adjusted p-value was corrected by the independent hypothesis weighting (IHW) method as implemented in the R package IHW v1.12 (Ignatiadis et al., 2016). Genes were considered differentially expressed when presenting an absolute log2 fold change equal or greater than 1.5 and an adjusted p-value lower than 0.05.

### In situ Hi-C

In situ Hi-C was performed as described (Rao et al., 2014) with slight alterations to the protocol. 5×10^6^ ESCs were crosslinked for 10 minutes in 1% formaldehyde before quenching in 125mM Glycine for 5 minutes. Cell pellets were washed twice in PBS before contact generation. Pellet was resuspended in 500uL Hi-C lysis buffer (10mM Tris-HCl pH8.0, 10mM NaCl, 0.2% NP40 and protease inhibitors) and incubated with rotation for 30 minutes at 4°C. Cells were pelleted and washed in 500uL Hi-C lysis buffer. Pellet was resuspended in 100uL 0.5% SDS and incubated for 10 minutes at 62°C. 285uL of H20 and 50uL of 10% Triton X-100 was added to quench the SDS at 37°C for 15 minutes. 50uL of 10xNEB DpnII buffer and 750U of DpnII were added (NEB R0453M) then incubated at 37°C overnight with shaking. Digestion was heat inactivated the next day for 20 minutes at 62°C. Biotin fill in was performed by addition of 52uL fill-in mix (0.288mM dCTP/dTTP/dGTP, 50U DNA Polymerase I, Large Klenow fragment (NEB M0210) and 0.288mM biotin-dATP (Thermo 19524016)). Fill-in reaction was incubated at 37°C for 1 hour with shaking. Ligation was performed by adding 150uL 10x NEB T4 DNA ligase buffer, 125uL 10% Triton X-100, 3uL 50mg/mL BSA and 8000U of T4 DNA ligase and H20 up to 1.5mL. Ligation was performed overnight at 25°C with shaking. Nuclei were pelleted before resuspending in IP buffer (33 mM Tris-HCl pH 8, 100 mM NaCl, 5 mM EDTA, 0.2% NaN_3_, 0.33% SDS, 1.66% Triton X-100) and sonicating to fragment size between 200-1000bp. 15uL of the sample was de-crosslinked overnight at 65°C with 85ul of decrosslinking buffer (0.1M NaHCO_3_, 1% SDS) before clean up using QiaQuick PCR clean up kit from Qiagen.

Library prep was performed as described (Mumbach et al., 2016). Streptavidin C-1 beads (Thermo 65001) were washed and resuspended in 2x biotin binding buffer (10mM Tris-HCl pH7.5, 1mM EDTA, 2M NaCl). 10ng of DNA was used for biotin capture, streptavidin C-1 beads were added and incubated for 15 minutes at room temperature with shaking. Beads were washed twice at 55°C for 2 minutes in tween wash buffer (5mM Tris-HCl pH7.5, 0.5mM EDTA, 1M NaCl, 0.05% Tween). Beads were washed once in 1xTD buffer (20mM Tris-HCl pH7.5, 10mM MgCl_2_, 20% DMF) before being tagmented with in-house produced Tn5 transposase in TD buffer for 55°C for 10 minutes. TN5 was quenched in 50mM EDTA for 30 minutes at 55°C before washing twice in 50mM EDTA, twice in Tween wash buffer and once in 10mM Tris. PCR amplification was carried out on beads using KAPA HotStart Taq enzyme. After purification of PCR product with Ampure beads, the quality of the obtained library was assessed by Bioanalyzer (High Sensitivity DNA kit, Agilent Technologies), prior to sequencing.

### HiC data analysis

DNA reads were filtered for low-quality bases and adapters using fastp (Chen et al., 2018). Filtered DNA reads were processed through the HiCpro pipeline to obtain contact matrices reporting the chromatin interactions at the genome-wide level (Servant et al., 2015). Using default settings, DNA reads were aligned to the mouse reference genome mm10 using bowtie2 in end-to-end mode, uniquely aligned DNA reads were assigned to the DpnII genomic restriction fragments, valid interactions were identified and used to generate interaction matrices. The HiCpro matrices containing all of the valid pairs were converted in *hic* format using the hicpro2juicebox function (Robinson et al., 2018), and in *cool* format using the hic2cool function (https://github.com/4dn-dcic/hic2cool). Contact matrices in cool format where corrected using the ICE algorithm as implemented in the hicCorrectMatrix function (Wolff et al., 2020). Plots of the interactions occurring at single locus or whole-chromosome level were obtained by HiCplotter (Akdemir and Chin, 2015).

### STORM imaging

For STORM imaging, 35mM MATTEK (P35G-1.5-14-C) plates were gelatinsed with 0.1% gelatin. 500,000 ESC were seeded per plate for 20-22 hours before 6 minute fixation in 1:1 Methanol: Ethanol at −20°C. Samples were blocked in blocking buffer (PBS with 10% donkey serum) for one hour at room temperature. Samples were incubated overnight at 4°C with H3 antibody (AB1791) 1:70 dilution in blocking buffer + 0.1% Triton X-100. Samples were incubated in 1:200 dilution of Alexafluor-647 secondary antibody in PBS in the dark for one hour at room temperature. Samples were then post-fixed in 2% PFA for 5 minutes before storing in PBS at 4°C until imaging.

Direct STORM (dSTORM) imaging was performed using the Nikon N-STORM microscope equipped with a 1.49 NA CFI Apochromat TIRF objective, exciting the Alexa Fluor 647 dye with the 647 nm laser light in HILO (highly inclined and laminated optical sheet) mode (Tokunaga et al., 2008). The 405 nm laser light (activation laser) was used for reactivating the Alexa Fluor 647 into a fluorescent state. The activation laser power was automatically increased by the NIS software to keep the number of localizations per frame constant up to a maximum of 50% of the laser power. Each dSTORM acquisition consisted of 40 thousand images recorded with an Orca-Flash4.0 sCMOS camera (Hamamatsu) with an exposure time of 20 ms, a pixel size of 161.5 nm and a field of view of 128×128 pixels. During dSTORM acquisitions, cells were kept in imaging buffer (100 mM MEA, 1% glucose, 560 ug/mL Glucose Oxidase and 34 ug/mL Catalase in PBS).

Two regions of interest (ROI) of 32×32 pixels for each dSTORM image were processed using ThunderSTORM (Ovesný et al., 2014) with a pre-detection wavelet filter (B-spline, scale 2, order 3), initial detection by non-maximum suppression (radius 1, threshold at one standard deviation of the F1 wavelet), and sub-pixel localization by integrated Gaussian point-spread function and maximum likelihood estimator with a fitting radius of 3 pixels. Detected localizations were filtered (intensity > 500 photons, sigma range of 50–500, and localization uncertainty < 20 nm). The filtered data set was then corrected for sample drift (cross-correlation of images from five bins at a magnification of 5) and repeated localizations was removed by merging points which reappeared within 20 nm. STORM images were visualized using the normalized Gaussian method with a lateral uncertainty of 20 nm.

Cluster analysis were performed thanks to a supervised machine learning approach using trained neural network (Williamson et al., 2020). The CAML (Cluster Analysis by Machine Learning) analysis workflow consisting of 3 stages and corresponding Python scripts, was used to: 1) prepare the data converting the x,y localization tables in a list of near-neighbour distances; 2) evaluate the input data with a trained model; 3) extract clustering information. The 87B144 model was used (Williamson et al., 2020), considering the possibility that more than 100 localizations per cluster could occur in our dataset.

### Mass spectrometry

For sub-cellular fractionation into cytosol, nucleosol and chromatin compartments, cell lysates were obtained according to established protocols (Méndez and Stillman, 2000). Briefly, to prepare nuclear extracts, the cells were washed twice with PBS and once with hypotonic buffer. Then, cells were incubated with hypotonic buffer (20 mM Hepes pH 8.0, 5 mM KCl, 1.5 mM MgCl_2_, 0.1 mM dithiothreitol [DTT]) for 30 min on ice. Triton X-100 (0.1%) was added, and the cells were incubated for 5 min on ice. Nuclei were collected in pellet by centrifugation (2 min, 3750 × *rpm*, 4°C). The supernatant was discarded and nuclei were washed once in hypotonic buffer, and then lysed in Urea buffer (8M Urea in 150 TrisHCl,pH 7.6, protease inhibitors). After 30 min on ice, insoluble proteins were removed from the nuclear extract by high-speed centrifugation (30 min, 13000 × *g*, 4°C). Protein extracts in UREA buffer were lysate, reduced, alkylated, digested and cleaned using the iST kit (Preomics).

For data analysis, the acquired tandem mass spectra were searched against the Uniprot mouse database using MaxQuant (version 1.6.2.3) (Tyanova et al., 2016a). Trypsin was specified as a cleavage enzyme, allowing up to three missed cleavages. Carbamidomethylation of cysteine was used as a fixed modification, with oxidation of methionine used as variable modifications. For label-free quantification the “match between runs” option in MaxQuant was applied. A false discovery rate of 1% was applied to peptide and protein identifications. Contaminants and reverse hits were excluded from the results prior to further analysis. Volcano plots were created using Perseus (version 1.6.2.3) (Tyanova et al., 2016b) (pValue=0.001).

## Data availability

Data were uploaded on the GEO repository with code GSEXXXXXX.

## Acknowledgments

We would like to acknowledge the IEO Genomic unit for support and all members of Pasini laboratory for helpful discussion. The work of the Pasini laboratory was supported by the Italian Association for Cancer Research, AIRC (IG-2017-20290) and by the European Research Council, ERC (EC-H2020-ERC-CoG-DissectPcG: 725268). E.C was supported by funding from AIRC and the European Union’s Horizon 2020 research and innovation programme under the Marie Sklodowska-Curie grant agreement (800924). D.F-P and EP are a PhD student within the European School of Molecular Medicine (SEMM). D.F-P was supported by a fellowship of FIRC and DM by a fellowship of the European Institute of Oncology Foundation (FIEO).

## Author contributions

E.C and D.P conceived the project. E.C performed the majority of the experimental work. F.R performed the majority of the computational analysis. S.T, E.P, K.J.F, M.Z and D.M provided support to the experimental work. D.F.-P. provided support to the computational analysis. S.T and E.P performed mass spectrometry experiments. S.R performed STORM imaging experiments and analysis. E.C and D.P wrote the manuscript and edited the figures.

## Declarations of Interests

The authors declare no competing interests.

## Notes

### Competing Interest Statement

The authors have declared no competing interest.

