## Supplementary Information for "BAP1 activity regulates PcG occupancy and global chromatin condensation counteracting diffuse PCGF3/5-dependent H2AK119ub1 deposition"

### Figure S1.

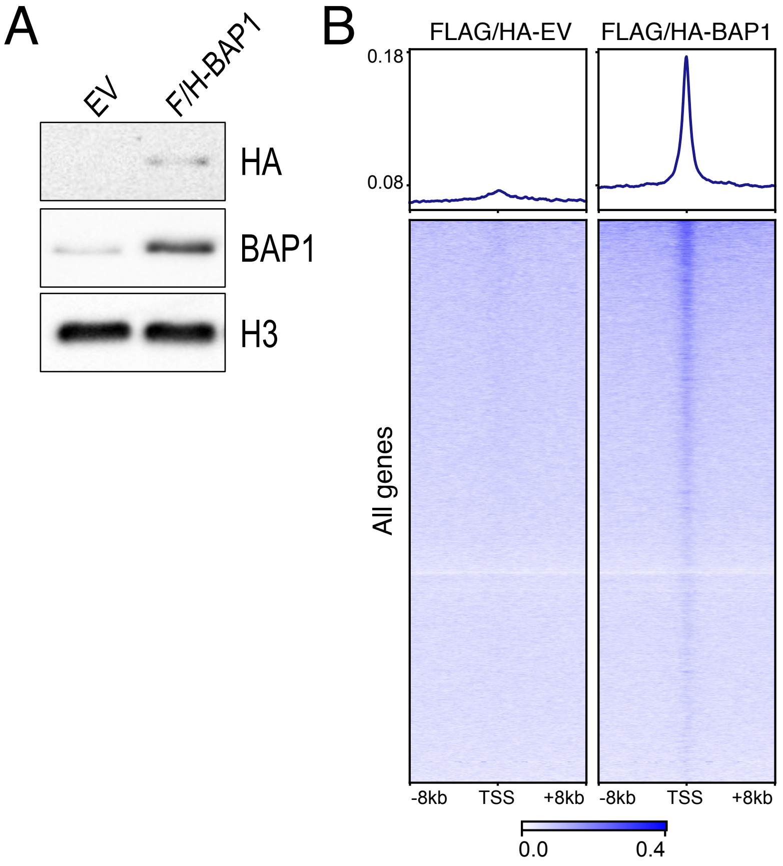
A
EV
F/H-BAP1
HA
BAP1
H3
B
FLAG/HA-EV
FLAG/HA-BAP1
0.18
0.08
All genes
-8kb
TSS
+8kb
-8kb
TSS
+8kb
0.0
0.4

**Figure S1. Supporting material for figure 1**

A) Western blot analysis with the indicated antibodies on total protein extracts from the indicated ESC cell lines (E14 WT + empty vector, E14 WT + FH-BAP1 WT).

**B)** Heatmaps representing normalized ChIP-seq intensity for HA in the E14 WT + empty vector and E14 WT + FH-BAP1 WT cell lines +/- 8kb of TSS. All TSS are shown.

### Figure S2.

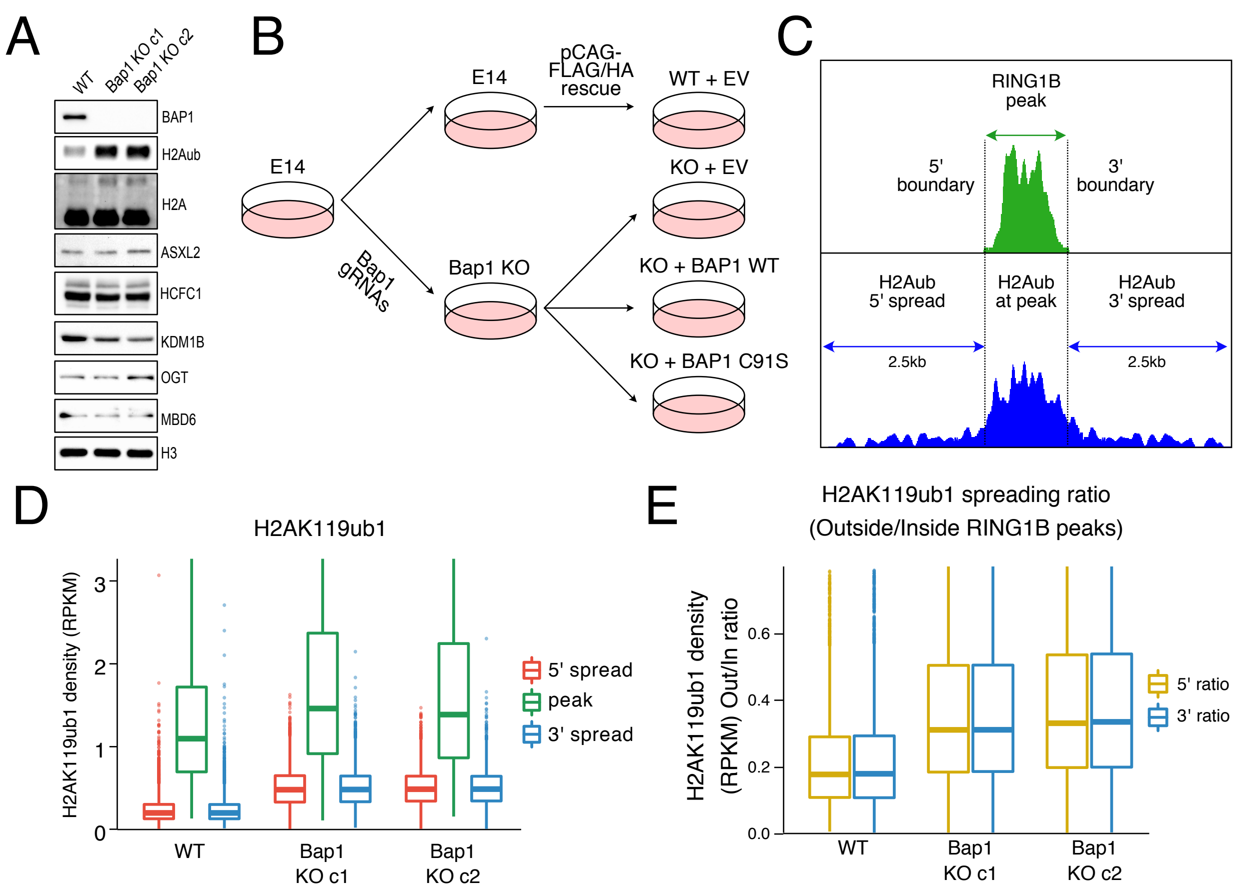

A
WT
Bap1 KO c1
Bap1 KO c2
BAP1
H2Aub
H2A
ASXL2
HCFC1
KDM1B
OGT
MBD6
H3
B
E14
E14
pCAG-FLAG/HA rescue
WT + EV
KO + EV
Bap1 gRNAs
Bap1 KO
KO + BAP1 WT
KO + BAP1 C91S
C
RING1B peak
5' boundary
3' boundary
H2Aub 5' spread
H2Aub at peak
H2Aub 3' spread
2.5kb
2.5kb
D
H2AK119ub1
H2AK119ub1 density (RPKM)
0
1
2
3
WT
Bap1 KO c1
Bap1 KO c2
5' spread
peak
3' spread
E
H2AK119ub1 spreading ratio
(Outside/Inside RING1B peaks)
H2AK119ub1 density (RPKM) Out/In ratio
0.0
0.2
0.4
0.6
WT
Bap1 KO c1
Bap1 KO c2
5' ratio
3' ratio

**Figure S2. Supporting material for figure 2**

A) Western blot analysis with the indicated antibodies on total protein extracts from the indicated ESC cell lines.

B) Schematic of strategy for generating BAP1 KO and either BAP1 WT or C91S mutant rescue cell lines.

C) Illustration describing criteria used to select RING1B inside peak regions, its boundaries, and the regions selected for the analysis at the 5' and 3' outside of RING1B peak boundaries.

D) Boxplots representing H2AK119ub1 density distributions in WT and BAP1 KO cells (two clones) within RING1B peaks, as well as 2.5kb outside of the 5' and 3' ends.

E) Boxplots representing the distribution of the H2AK119ub1 density ratio between the H2AK119ub1 density inside RING1B peaks and at 5' or 3' spreading regions

### Figure S3.

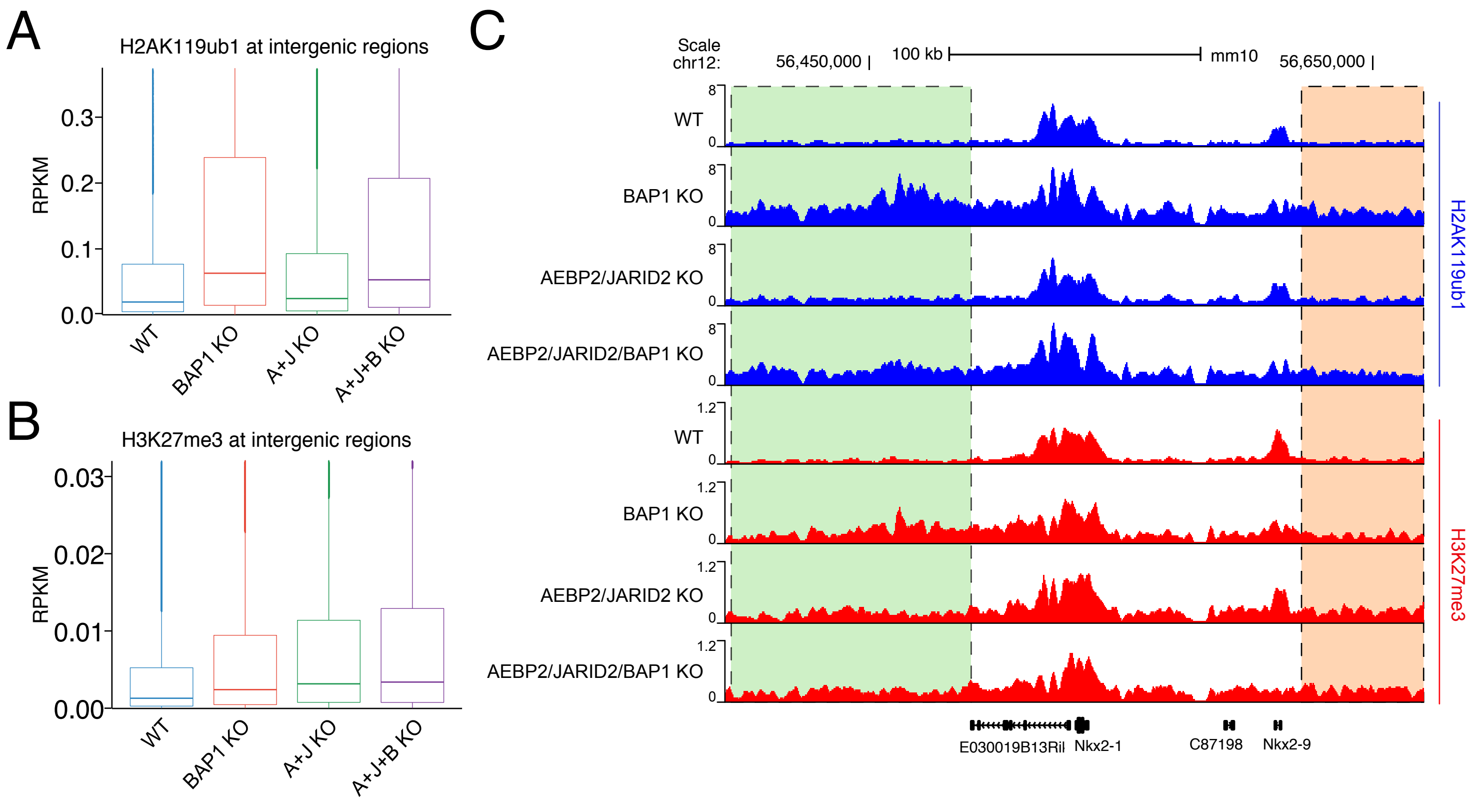
A
H2AK119ub1 at intergenic regions
RPKM
0.0
0.1
0.2
0.3
WT
BAP1 KO
A+J KO
A+J+B KO
B
H3K27me3 at intergenic regions
RPKM
0.00
0.01
0.02
0.03
WT
BAP1 KO
A+J KO
A+J+B KO
C
Scale
chr12:
56,450,000 |
100 kb
mm10
56,650,000 |
WT
BAP1 KO
AEBP2/JARID2 KO
AEBP2/JARID2/BAP1 KO
WT
BAP1 KO
AEBP2/JARID2 KO
AEBP2/JARID2/BAP1 KO
8
0
1.2
0
H2AK119ub1
H3K27me3
E030019B13Ril
Nkx2-1
C87198
Nkx2-9

**Figure S3. Supporting material for figure 5**

A) Boxplots representing H2AK119ub1 ChIP-seq RPKM levels in the indicated cell lines at intergenic sites.
B) Boxplots representing H3K27me3 ChIP-seq RPKM levels in the indicated cell lines at intergenic sites.
C) UCSC genome browser snapshot of H2AK119ub1 and H3K27me3 ChIP-seq in the indicated cell lines.

### Figure S4.

A

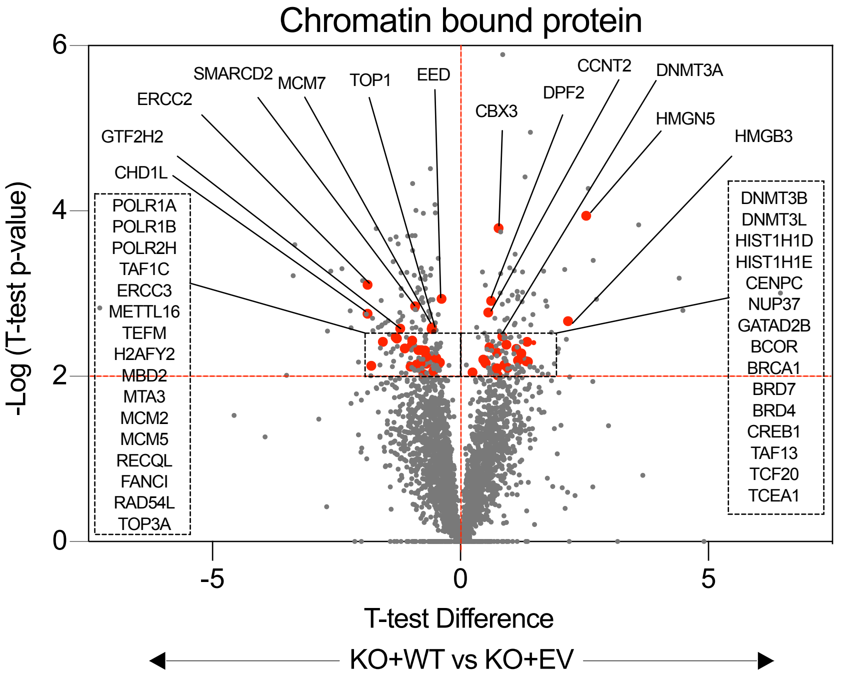

Chromatin bound protein
-Log (T-test p-value)
T-test Difference
KO+WT vs KO+EV
6
4
2
0
-5
0
5
SMARCD2
MCM7
TOP1
EED
ERCC2
GTF2H2
CHD1L
CBX3
DPF2
CCNT2
DNMT3A
HMGN5
HMGB3
POLR1A
POLR1B
POLR2H
TAF1C
ERCC3
METTL16
TEFM
H2AFY2
MBD2
MTA3
MCM2
MCM5
RECQL
FANCI
RAD54L
TOP3A
DNMT3B
DNMT3L
HIST1H1D
HIST1H1E
CENPC
NUP37
GATAD2B
BCOR
BRCA1
BRD7
BRD4
CREB1
TAF13
TCF20
TCEA1


B

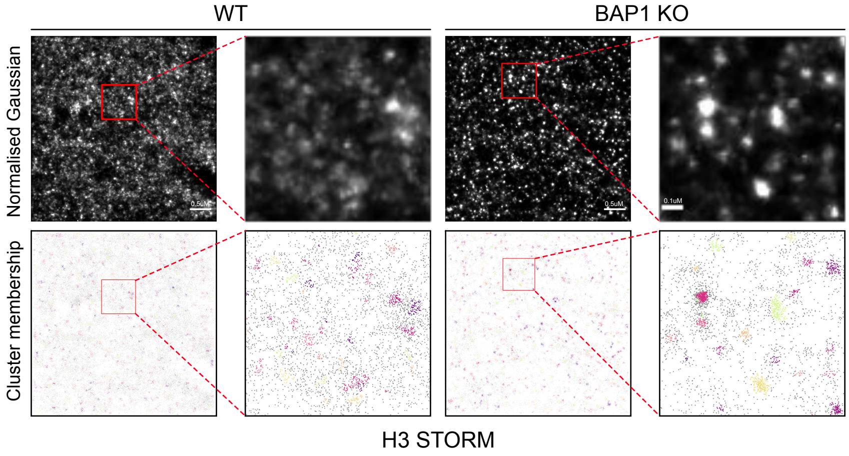

WT
BAP1 KO
Normalised Gaussian
Cluster membership
H3 STORM

**Figure S4. Supporting material for figure 6**

A) Volcano plot of mass spectrometric analysis of chromatin bound proteins in BAP1 KO+BAP1 WT vs BAP1 KO+EV. pValue=0.001

B) Representative STORM images of WT and BAP1 KO ESC stained with Histone H3 antibody. Both normalised gaussian and cluster membership images are shown for the same regions of interest.
